# Small-molecule ACOD1 inhibition attenuates colitis and tumor growth through a shared myeloid mechanism, diverging from genetic deletion

**DOI:** 10.64898/2026.10.17.759827

**Authors:** Adonia E. Papathanassiu, Weixing Li, Konstantin Gilbo, Ian L. Pegg

**Affiliations:** Ergon Pharmaceuticals, LLC, P.O. Box 1001, Silver Spring, MD 20910, USA; Vitreous State Laboratory, The Catholic University of America, Washington, DC 20064, USA

## Abstract

Aconitate decarboxylase 1 (ACOD1), the enzyme responsible for synthesis of the immunometabolite itaconate, is widely regarded as an anti-inflammatory brake on tissue damage based on *Acod1*-deficient mice and itaconate supplementation expermiments. ERG344, a small-molecule ACOD1 inhibitor, suppressed itaconate synthesis in endotoxemic mice and in inflamed colonic mucosa and, contrary to genetic-deletion studies, attenuated acute DSS colitis in an ACOD1-dependent manner. ERG344 treatment was associated with reduced influx of immature monocytes early on and accumulation of mature neutrophils. In syngeneic tumor models, ERG344 prolonged survival and increased tumor-infiltrating lymphocytes and antigen-presenting dendritic cells. Across murine and human colitis scRNA-seq datasets and a ten-model syngeneic tumor atlas, ACOD1 marked interferon-stimulated immature monocytes and maturing neutrophils, and tracked with *SOD2* and *CD274* independently of interferon and NF-κB signaling. ACOD1 appears to act through a common myeloid recruitment and maturation mechanism, making pharmacological ACOD1 inhibition a viable therapeutic strategy in colitis and cancer.

## Introduction

Aconitate decarboxylase 1 (ACOD1, formerly IRG1) is an inducible mitochondrial enzyme, primarily known for diverting the tricarboxylic acid (TCA) cycle of inflammatory myeloid cells towards production of itaconate through decarboxylation of *cis*-aconitate (1). Itaconate is a powerful immunometabolite that modulates inflammation through multiple and often parallel mechanisms (2). As a structural analog of succinate, it competitively inhibits succinate dehydrogenase (SDH) thereby decreasing ROS production, HIF-1α stabilization, and downstream pro-IL-1β transcription (3). Its electrophilic α,β-unsaturated dicarboxylate core enables post-translational modification of protein cysteine residues via electrophilic alkylation. Key alkylation targets include KEAP1 (which releases NRF2 to drive an antioxidant transcriptional program) (4), the IκBζ-ATF3 axis (which selectively restrains IL-6 and IL-12 transcription) (5), NLRP3 (which blocks inflammasome assembly) (6), and rate-limiting glycolytic enzymes (which suppress the metabolic rewiring required for macrophage activation) (2). Beyond host pathways, itaconate exerts direct antimicrobial activity by inhibiting bacterial isocitrate lyase (2). Its functional consequences are considered strictly context-dependent. Despite its anti-inflammatory properties, unmodified itaconic acid can also enhance type I interferon signaling, while specific pathogens can co-opt its electrophilicity to evade host clearance (2). Adding to this complexity, recent studies have uncovered entirely itaconate-independent functions of ACOD1, such as the induction of CD274/PD-L1 expression in proinflammatory monocytes and the direct inhibition of CD8^+^ T cell proliferation (7, 8). As a result, the current consensus is that the impact of ACOD1 in a specific disease does not depend on a common mechanism of action but rather on the cellular and metabolic context of the disease, which determines which itaconate function predominates.

ACOD1 expression has been reported in numerous pathologies including bacterial and viral infections (9–11), in ischemic heart disease (12), in atherosclerotic cardiovascular disease (13), and various cancers (14–16). In inflammatory bowel disease (IBD), *ACOD1* gene expression has been reported in the inflamed mucosa of patients with Crohn’s disease (CD) and ulcerative colitis (UC), where it positively correlates with the intensity of intestinal inflammation (17,18). It is generally believed that ACOD1 is upregulated at sites of severe inflammation to produce itaconate and to limit colonic tissue damage (19). This view stems from studies demonstrating that *Acod1* deficiency in mice exacerbates the clinical symptoms and histological severity of dextran sulfate sodium (DSS)-induced colitis and confirmed in part by studies showing mitigation of the disease in wild-type (wt) mice through treatment with a cell-permeable itaconate-derivative such as 4-octyl itaconate (4-OI) (18, 20–22). Although ACOD1 expression has typically been associated with pro-inflammatory macrophages (1), single cell RNA sequencing (scRNA-seq) studies in DSS colitis have suggested that the predominant source of *Acod1* in intestinal inflammation is mature activated neutrophils and a distinct, hyper-inflammatory subpopulation of monocytes with colonic macrophages characterized by the absence of *Acod1* (18,22,23). In a preprint publication, Jones et al. reported that, in human IBD, *ACOD1* presence was restricted to newly recruited monocytes, which acquired the gene shortly after arrival at the inflamed colon and that, histologically, ACOD1 marked sites of extensive mucosal damage (18). Additionally, Zhao et al. linked *Acod1* deletion with an altered hematopoiesis, increased neutrophil infiltration into the colon, upregulation of inflammation, and enhanced reverse neutrophil migration (22). Collectively, the above studies point to a critical spatiotemporal regulation of ACOD1 that goes beyond a generalized anti-inflammatory brake.

In contrast to colitis, where ACOD1 induction appeared to be host-protective, ACOD1 expression in cancer has been associated with the rise of an immunosuppressive microenvironment and immune evasion of tumor cells (14–16). Peritoneal tissue-resident macrophages from tumor-bearing animals upregulated Acod1 and accumulated itaconate over 15-fold to promote peritoneal melanoma and ovarian tumor growth (24). Malignant cells facilitated induction of Acod1 in tumor-associated macrophages (TAMs) via a NF-κB mechanism, while TAM-produced itaconate epigenetically silenced *Cxcl9* and *Cxcl10* expression through inhibition of DNA dioxygenase TET2 and restricted antigen presentation and T cell migration into the tumor core (15). In tumor-bearing immunocompetent mice, *Acod1* deletion was associated with increased monocyte infiltration, a higher frequency of pro-inflammatory TAMs, and an elevated abundance of tumor-infiltrating lymphocytes (TILs) (15). *Acod1*^−/−^ mice survived significantly longer than their wt counterparts across diverse tumor challenges, with *Acod1* deficiency synergizing with anti-PD-L1 checkpoint blockade to further improve survival (15). While *Acod1* was also expressed by tumor-associated neutrophils (TANs), the latter had no measurable impact on tumor growth, as demonstrated by neutrophil depletion experiments (15). Overall, these findings establish ACOD1 as a potent immunosuppressive driver within the tumor microenvironment, whose disruption invigorates anti-tumor immunity and enhances the efficacy of both checkpoint inhibitors and myeloid cell-based therapies. However, as in the case of colitis, the view of the role of ACOD1 in cancer has been shaped almost exclusively by germline deletion studies. Because constitutive gene deletion can induce compensatory adaptations with unforeseen functional consequences, we sought to examine the effects of pharmacological inhibition of ACOD1 in DSS colitis and cancer using a novel small-molecule inhibitor, ERG344. Our results challenge two conventions regarding ACOD1: the idea that its functions are context-dependent and that ACOD1 has a net beneficial effect in DSS colitis.

## Results

### Characterization of ERG344

Small-molecule inhibitor ERG344 (disodium 2-benzyl-3-[(4-fluorophenyl)methylidene]butanedioate) was designed around the succinate backbone of itaconate (Fig. 1A) and synthesized as a salt to ensure aqueous solubility. Its ability to inhibit ACOD1 was evaluated by measuring the amount of itaconate, produced by rat C6 glioma cells that have been engineered to overexpress ACOD1. ACOD1 produces itaconate per the reaction shown in Supplementary Fig. S1A, typically as part of the inflammatory TCA cycle (Supplementary Fig. S1B). In Acod1^+^ C6 cells, treatment with 200 μM ERG344 significantly reduced intracellular itaconate levels to ∼45% of vehicle-treated controls (Fig. 1B), confirming cellular target engagement. Pharmacokinetic (PK) evaluation of ERG344 in C57BL/6 mice demonstrated favorable drug-like properties. Intravenous (i.v.) administration of 1 mg/kg (mpk) ERG344 led to an initial concentration (C_0_) of ∼60 μM and an elimination half-life (t_1/2_) of ∼8.3h as shown in Supplementary Fig. S1C. Oral administration of 5 mpk ERG344 resulted in rapid systemic absorption (T_max_=0.5h) and high oral bioavailability (F∼73%; Supplementary Fig. S1C). Because molecules with the itaconate backbone structure can activate the cytoprotective NRF2 pathway, which has been shown to be responsible for many of itaconate’s anti-inflammatory properties, we examined whether ERG344 directly alters NRF2 signaling. Exposure of THP-1 monocytes to ERG344 stabilized NRF2 protein levels but induction was concentration-dependent (Fig. 1C). While no significant activation occurred at 125 µM, a twofold increase in NRF2 protein levels was observed at 500 µM ERG344 (Fig. 1C). Based on these findings and assuming linear scaling, i.v. doses of ≥8 mpk ERG344 are projected to reach drug concentrations sufficient to activate NRF2, while lower doses should leave the pathway unaffected. Notably, the majority of in vivo studies described here utilized dosing of ERG344 at ≤5 mpk, given intraperitoneally (i.p.) once or twice a day. The pharmacological properties of Supplementary Fig. S1C suggest that a dose of 5 mpk ERG344 will result in a steady state concentration of ∼80-120 μM. Furthermore, ERG344 exhibits a low V_ss_ value, which argues against tissue accumulation. The data collectively indicate that, at doses ≤5 mpk, ERG344 is unlikely to activate the NRF2 pathway.

**Figure 1.**
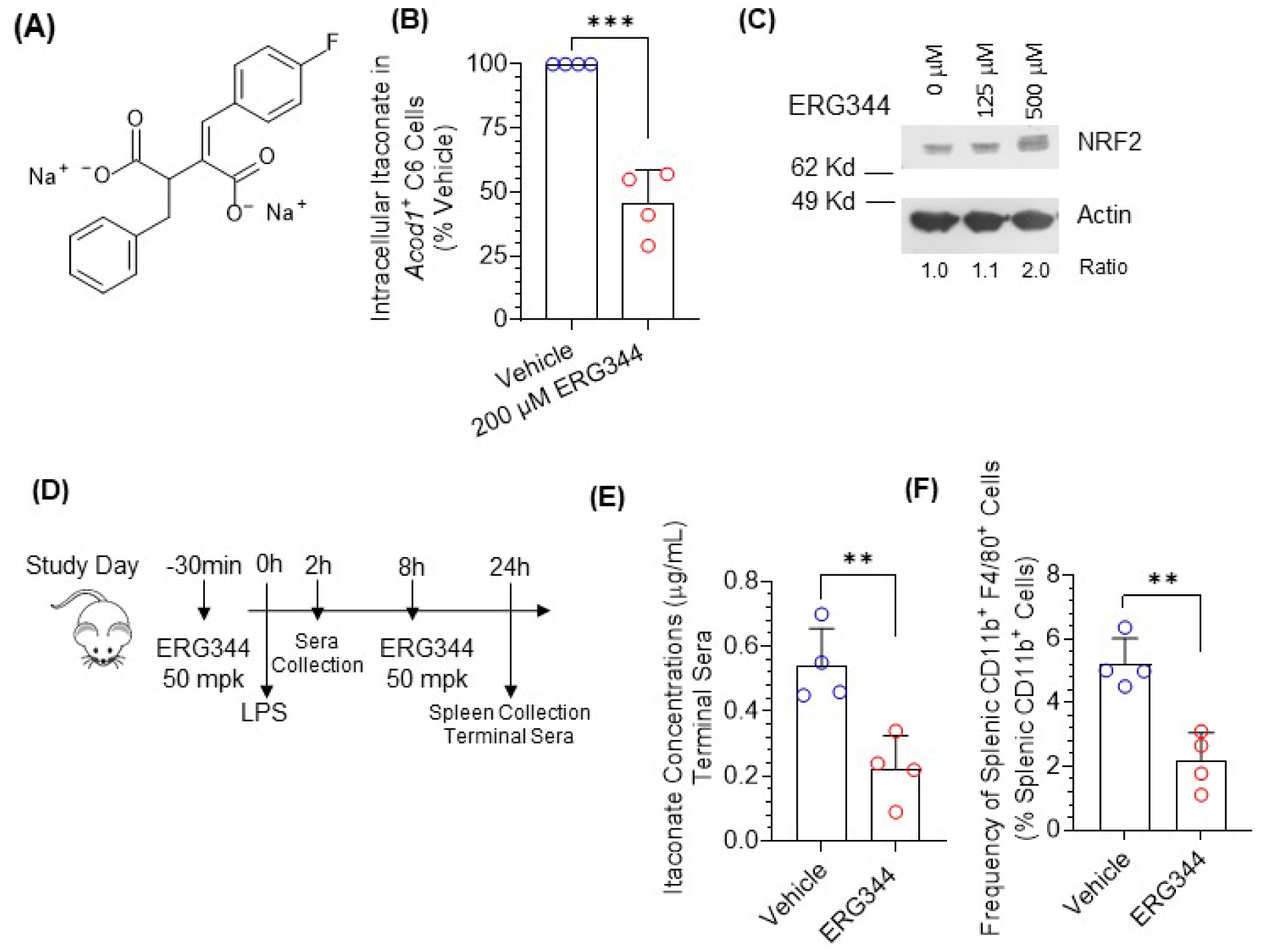
ERG344 inhibits itaconate synthesis in vitro and in vivo. (A) Structure of ERG344 (disodium 2-benzyl-3-[(4-fluorophenyl)methylidene]butanedioate). (B) Intracellular itaconate in Acod1^+^ rat C6 glioma cells treated overnight with vehicle or 200 μM ERG344, expressed as percentage of vehicle (n=4 independent experiments). (C) Western immunoblotting showing the effect of various concentrations of ERG344 on NRF2 expression in THP-1 cell lysates, collected at 4h of treatment. (D) Design of the acute LPS endotoxemia study: ERG344 (50 mpk, i.p.) 30 min before and 8 h after LPS challenge; serum was collected at 2 and 24 h, spleens and terminal sera at 24 h. (E) Serum itaconate at 24h of LPS challenge (n=4 mice/group). (F) Frequency of splenic CD11b⁺F4/80⁺ macrophages among splenic CD11b⁺ cells at 24 h. Data are mean±SD; **P<0.01; *** P<0.001.

We next evaluated the ability of ERG344 to suppress itaconate production in vivo in a mouse model of acute lipopolysaccharide (LPS)-induced endotoxemia (Fig. 1D). Administration of 50 mpk ERG344 i.p. significantly blunted circulating itaconate levels, reducing serum concentrations from ∼0.5 μg/mL in vehicle-treated controls to ∼0.2 μg/mL, a ∼60% reduction (Fig. 1E). This metabolic blockade correlated with a significant shift in systemic myeloid composition as the frequency of CD11b^+^F4/80^+^ macrophages in the spleen was reduced after ERG344 treatment (Fig. 1F). It is well-understood that mice challenged with LPS respond with a rapid increase in circulating TNFα and that itaconate alleviates TNFα production (25,26). However, no significant changes in the concentration of circulating TNFα were seen here with ERG344 treatment (Supplementary Fig. S1D), although the lack of effect might be a timing issue.

### ERG344 protects against DSS-induced colitis in an ACOD1-dependent manner

To evaluate the impact of systemic ACOD1 inhibition on colonic inflammation, we administered ERG344 (5 mpk, ip) to wt C57BL/6 mice receiving 2% DSS in their drinking water, beginning 24h after initiation of the DSS challenge. We monitored the animals for signs of disease for seven days (Fig. 2A). Acute DSS colitis follows a predictable trajectory. Epithelial barrier breakdown is recorded within 24 h (with bacteria crossing the inner mucus layers within 4-12h), increased permeability is seen by day 3, inflammatory marker induction by day 4, onset of weight loss by day 5, and peak disease severity, characterized by rectal bleeding, severe weight loss, and extensive mucosal damage, by day 7 (27). In Fig. 2B, ERG344 treatment reduced colonic lamina propria (LP) itaconate levels to ∼30% of vehicle control values by day 7 of DSS administration, confirming target engagement within the inflamed gut mucosa, and attenuated the clinical severity of colitis. ERG344-treated wt mice exhibited a lower Disease Activity Index (DAI) from day 5 onward (with a reduction of ∼50% in terminal DAI compared to vehicle controls) (Fig. 2C) and were protected against progressive body weight loss (Fig. 2D). Moreover, ERG344 largely prevented the colitis-induced increase in colon mass-to-length ratio (a metric of tissue edema, wall thickening, immune cell infiltration, fibrosis, and colon shortening), maintaining it at baseline levels (Fig. 2E). Because the findings directly contradict published reports showing that genetic *Acod1* deficiency exacerbates colitis, we sought to confirm target specificity by subjecting *Acod1*^−/−^ mice to an identical DSS challenge and ERG344 treatment (Figs. 2F–2H). The protective effects of ERG344 were largely abrogated in *Acod1*^−/−^ mice with DAI scores and weight loss trajectories being statistically indistinguishable between vehicle-and ERG344-treated animals by day 7 (Figs. 2F–2G). Likewise, while there was a minor trend toward protection against colon shortening, this difference was not statistically significant, and ERG344 failed to prevent the increase in colonic mass-to-length ratio in *Acod1*^−/−^ mice (Fig. 2H). Images of resected colons from the experiments with wt and Acod1^-/-^ mice can be found in Supplementary Figs. S2A and S2B, respectively. Notably, because the wt and *Acod1*^-/-^ mice were obtained from different vendors and have different substrain backgrounds, Fig. 2 only assesses the effects of ERG344 on colitis progression within the same substrain. Collectively, the above data suggest that ERG344 protects against acute colonic inflammation primarily through an on-target, ACOD1-dependent mechanism.

**Figure 2.**
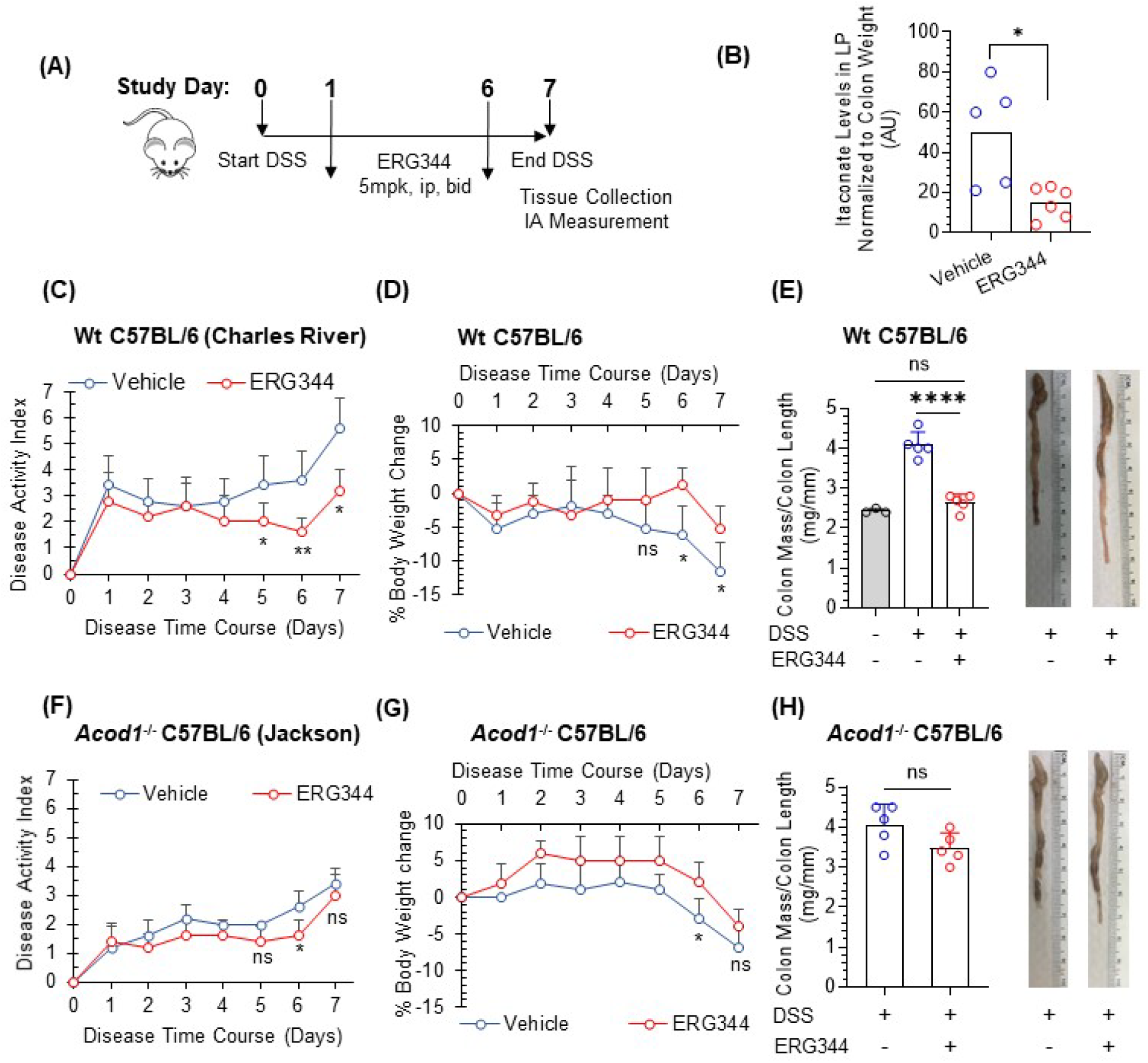
ERG344 attenuates DSS colitis through an ACOD1-dependent mechanism. (A) Study design of an acute DSS colitis model. (B-E) Data from wt C57BL/6 mice. (B) Itaconate present in colonic tissue lysates on day 7, normalized to colon weight (arbitrary units; n=5 mice/group; ERG344 group includes a technical replicate). (C) Disease activity index (DAI). (D) Body weight change from day 0. (E) Colon mass-to-length ratio on day 7 with representative resected colons. Ratios from DSS-naive animals are historic values shown for reference. (F–H) Data from *Acod1*^⁻/⁻^ C57BL/6 mice on an identical protocol (n=5 mice/group) (F) DAI. (G) Body weight change from day 0. (H) Colon mass-to-length ratio with representative colons. Data are mean± SD. ns: not significant; *P<0.05; **P<0.01; ****P<0.0001.

### ACOD1 blockade limits immature myeloid cell infiltration into the inflamed mucosa

Normal immune cell development in the gut follows a well-described continuum (also known as “monocyte waterfall”) from newly recruited blood monocytes to terminally differentiated resident macrophages. This maturation can be tracked via flow cytometry using specific surface markers, including CX3CR1, MHCII (indicating acquired antigen-presenting capacity), and CD64 (the canonical resident macrophage marker) (28). Subsequently, we evaluated whether ERG344 alters LP immune populations during early monocyte influx (day 4; Fig. 3A) and peak inflammatory infiltration (day 7; Fig. 3D) in acute DSS colitis. Flow cytometric analysis revealed that by day 4, ERG344-treated mice exhibited a reduction in total LP myeloid (CD11b^+^CD11c^-^) cells (Fig. 3B). This cell compartment was further depleted of freshly recruited, immature (CD11b^+^CD11c^-^ CX3CR1^lo/int^CD64^-^MHCII^-^) monocytes, whose frequency within myeloid cells was reduced from ∼32% in vehicle control mice to ∼6% following ERG344 treatment (Fig. 3C). Notably, ERG344 spared transitioning (CD11b^+^CD11c^-^CX3CR1^int^CD64^-^MHCII^+^) monocytes (Supplementary Fig. S2C) and mature resident (CD11b^+^CD11c^-^CX3CR1^hi^CD64^+^) macrophages (Supplementary Fig. S2D). Together, the data indicate that ERG344 specifically limits the mucosal influx of new, immature monocytes rather than depleting differentiating or established resident macrophage pools. Parallel transcriptomic analysis of colonic tissues via qRT-PCR revealed no significant differences in the expression of major inflammatory genes, *Acod1*, or *Hmox1* (Supplementary Fig. S2E). Yet, ERG344-treated mice maintained colon mass-to-length ratios at baseline levels, lower than those of vehicle controls (Supplementary Fig. S2F). We attributed the protective effect of ERG344 to the lower number of freshly recruited monocytes reaching the colon rather than on an effect on inflammatory pathways.

**Figure 3.**
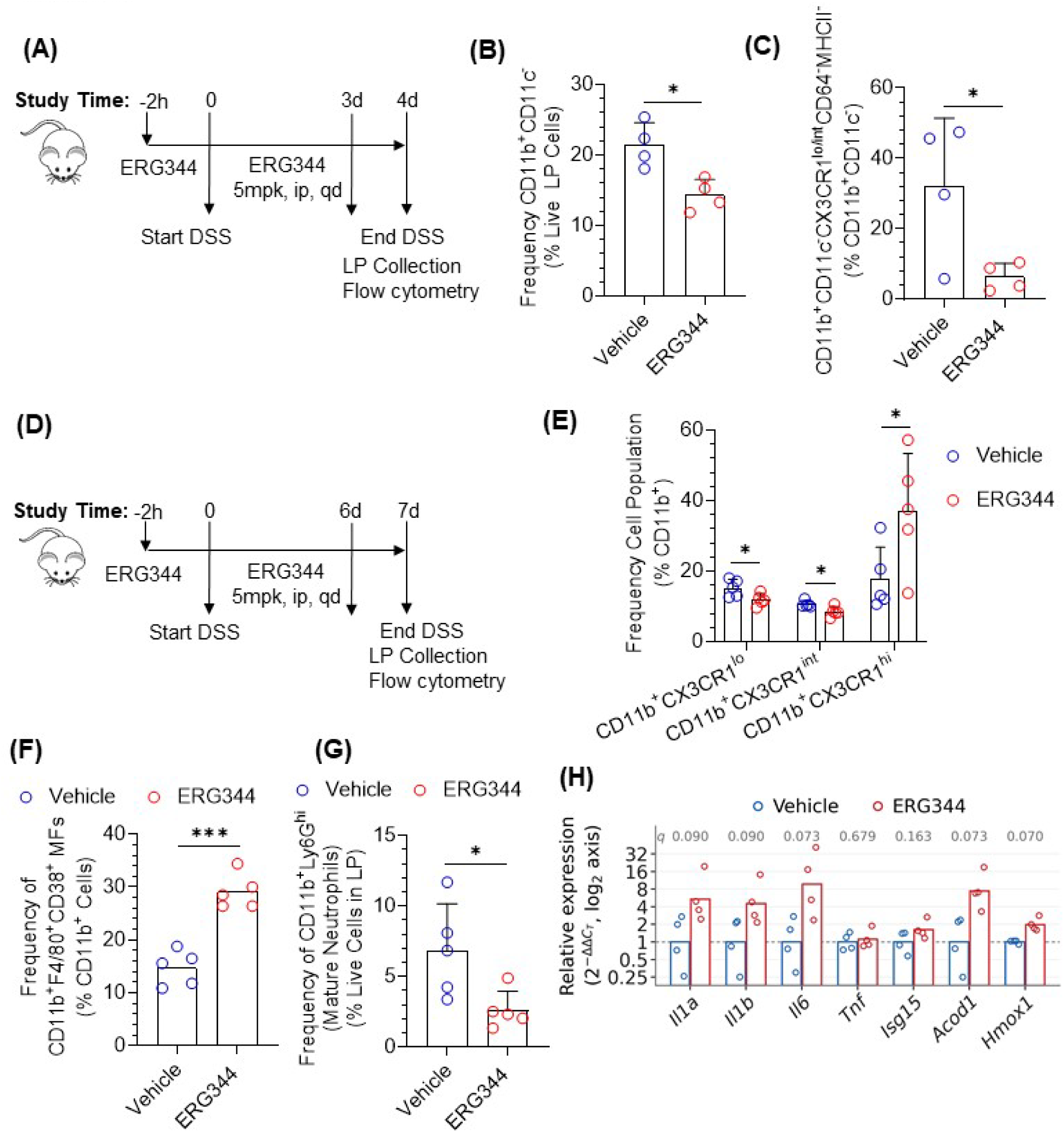
ERG344 restricts early immature monocyte influx and blunts peak neutrophil accumulation in the inflamed mucosa. (A) Study design of the early inflammation DSS colitis. ERG344 (5 mpk, i.p., qd). LP cells were collected on day 4 (n=4 mice/group). (B) Frequency of CD11b^+^CD11c^-^ myeloid cells among live LP cells on day 4. (C) Frequency of immature (CD11b^+^CD11c^-^CX3CR1^lo/int^ CD64^-^MHCII^-^) monocytes among LP CD11b^+^CD11c^-^ cells on day 4. (D) Study design of the peak-inflammation DSS colitis. ERG344 (5 mpk, i.p., qd). LP cells were collected on day 7 (n=5 mice/group). (E) Frequencies of CD11b^+^CX3CR1^lo^, CX3CR1^int^ and CX3CR1^hi^ cells among LP CD11b^+^ cells on day 7. (F) Frequency of pro-inflammatory CD11b^+^F4/80^+^CD38^+^ macrophages among LP CD11b^+^ cells on day 7. (G) Frequency of mature neutrophils (CD11b^+^Ly6G^hi^) among live LP cells on day 7. (H) qRT-PCR of colonic tissues on day 7, normalized to *Actb*. FDR q values are shown in the graph. Data are mean±SD. ns: not significant; *P<0.05; ***P<0.001.

A parallel analysis of colonic tissues on day 7 of DSS administration revealed that the population of immature (CD11b^+^CX3CR1^lo/int^) monocytes in ERG344-treated mice had largely rebounded, although it remained slightly below that of vehicle controls (Fig. 3E). An increase in the frequency of mature resident macrophages was also noted in the ERG344 cohort (Fig. 3E).

Consistent with the depletion of immunosuppressive itaconate by ERG344, the frequency of pro-inflammatory (CD11b^+^F4/80^+^CD38^+^) macrophages in the LP markedly increased from ∼14% to ∼28% of the total myeloid compartment (Fig. 3F). Concurrently, the frequency of mature neutrophils (CD11b^+^Ly6G^hi^) was reduced by >60% (Fig. 3G). Finally, transcriptomic analysis of colonic tissues at day 7 revealed a trend of increased inflammation (with increasing levels of *Il1a*, *Il1b*, *Il6* but not *Tnf or Isg15)* along with elevated amounts of *Acod1*, possibly as a feedback mechanism (Fig. 3H).

### Analysis of ACOD1 expression in human and murine colitis by scRNA-seq

To understand the possible effect of ACOD1 on myeloid production and/or recruitment, we investigated *Acod1* dynamics during intestinal inflammation by reanalyzing publicly available scRNA-seq datasets. We first focused on a longitudinal study of murine DSS colitis published by Ho et al. in 2021 (29), which examined colonic immune cell populations over a 15-day period (sampled at days 0, 3, 6, 9, 12, and 15) after challenging mice with 1.5% DSS in their drinking water for the first 6 days, followed by a recovery phase. The original investigators analyzed macrophages (classified as “mononuclear phagocytes” or MNPs) and neutrophils (classified as “granulocytes”) broadly, reporting that macrophage infiltration peaked at day 6 and normalized by day 15, whereas neutrophil accumulation peaked at day 9 and remained elevated through the end of the study. Reanalyzing this dataset for the presence of *Acod1*, we determined that *Acod1* expression was myeloid-restricted, appearing predominantly in neutrophils (92.6%) and secondarily in MNPs (15.6%) (Fig. 4A). The finding was in agreement with previously reported observations (18,22). We then independently re-clustered MNP cells (Fig. 4B) and annotated them using the framework suggested by Kang et al. (30). We identified six distinct subpopulations (Supplementary Fig. S3A). Within these MNP clusters, *Acod1* expression was predominantly associated with the ISG-Mono cluster, the monocyte subpopulation that highly expressed interferon-stimulated genes (ISGs). In that population, *Acod1* expression tracked with disease progression, peaking at day 6 and again at day 15 at the end of the monitored recovery time (Fig. 4C). At day 6 of disease, >60% of ISG-Monocytes and nearly 20% of transitional MoDMs expressed *Acod1*. Moreover, the expression of *Acod1* was correlated with ISGs (Supplementary Fig. S3B), confirming previous reports characterizing *Acod1* as an interferon-inducible gene (31). To define the transcriptional state marked by *Acod1*, we compared *Acod1*^+^ (n=482) and *Acod1*^-^ (n=1,412) monocytes comprising the Ly6C^hi^, ISG-Mono, and Trans-MoDM clusters. The analysis revealed that *Acod1* expression is linked to the concurrent upregulation of both pro- and anti-inflammatory transcriptional programs. Induction of some pro-survival and canonical NRF2 target genes, which can be activated by itaconate, was also observed (Supplementary Fig. S3C). Further investigation revealed that the majority of upregulated genes in *Acod1*^+^ monocytes were NF-κB inducible (Supplementary Fig. S3D). Collectively, the above analyses suggest that *Acod1* expression identifies a distinct, hyper-inflammatory monocyte state characterized by robust activation of both interferon and NF-κB signaling pathways, an observation that aligns with previously reported findings (18).

**Figure 4.**
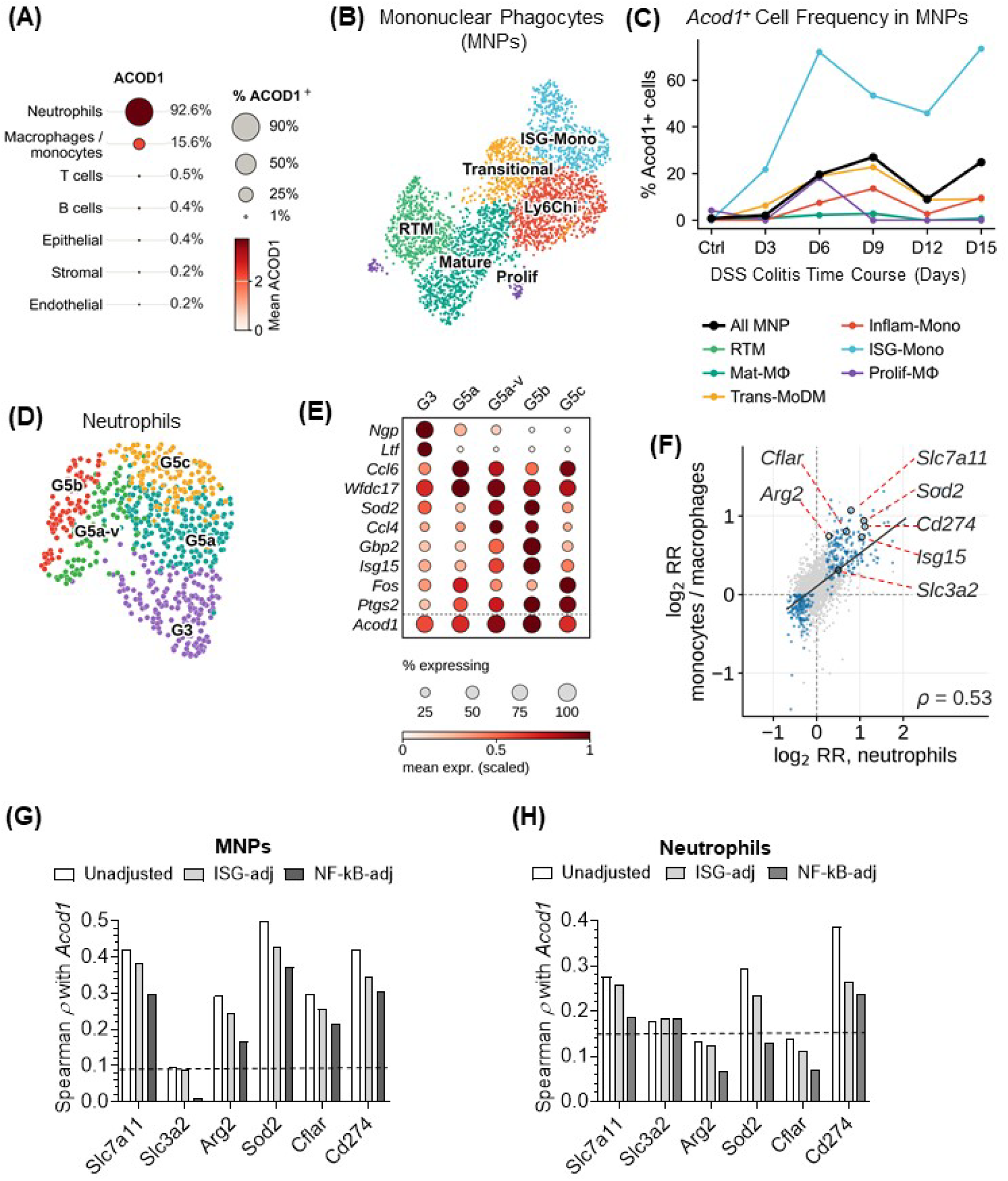
*Acod1* marks interferon-stimulated immature monocytes and maturing neutrophils in DSS colitis. Reanalysis of the longitudinal DSS colitis scRNA-seq dataset of Ho et al. (29): 1.5% DSS for 6 days followed by recovery, sampled on days 0, 3, 6, 9, 12 and 15. (A) Dot plot of *Acod1* across major colonic lineages; dot size, percentage of cells with detected transcript; color: mean scaled expression. (B) UMAP of re-clustered mononuclear phagocytes (MNPs) annotated using the framework of Kang et al. (30). The following clusters were identified: a) resident tissue macrophages (RTMs), representing self-maintaining, embryo-derived tissue-resident macrophages; b) mature monocyte-derived macrophages (MoDMs), corresponding to the P3 stage according to Bain et al. (28), c) transitional MoDMs, an intermediate population of differentiating macrophages corresponding to the P2 stage, d) inflammatory monocytes, corresponding to freshly recruited Ly6C^hi^ monocytes at the P1 stage, e) monocytes with high expression of interferon-stimulated genes (ISG-Mono), and f) a small cluster of proliferating macrophages (Prolif). (C) Percentage of *Acod1^+^* cells within each MNP subpopulation across the disease time course. (D) UMAP of re-clustered colonic neutrophils, resolved into one immature (G3) and four mature (G5a, G5a-v, G5b, G5c) states. (E) Dot plot of maturation and activation markers plus *Acod1* across neutrophil states. (F) Concordance of *Acod1* co-expression between neutrophils (x) and MNPs (y). Per-gene rate ratios (log₂, median-centred) from separate depth- and day-adjusted models. Blue: FDR<0.05 in both. Spearman ρ = 0.53. Genes of interest as labeled. (G, H) Spearman correlation of six selected genes with *Acod1* in (G) MNPs and (H) neutrophils, unadjusted and after adjustment for interferon (ISG-adj) or NF-κB (NF-κB-adj) activity. Dashed line, 95th percentile of the permutation null obtained by shuffling *Acod1* expression.

We also re-clustered neutrophils present in the Ho et al dataset (29). The cells were resolved into five distinct states: one immature (G3) and four mature ones (G5a, G5a-variant, G5b, and G5c) (Figs. 4D–4E). Unlike the MNP compartment, *Acod1* was ubiquitously expressed across >90% of all colonic neutrophils operating as a continuous gradient rather than a binary marker. Its expression peaked specifically within the interferon-activated states (G5b and G5a-variant), mirroring what was seen in monocytes. We then tested the transcriptome for genes whose expression rises or falls with *Acod1*, fitting per-gene Poisson rate models separately in neutrophils and MNPs with log total counts as an offset and adjustment for batch and disease day. Genes associated with *Acod1* at FDR<0.05 in both population formed a shared signature of 319 genes (Fig. 4F), from which we selected six for further evaluation. These genes included a cystine/redox group (*Slc7a11*, *Slc3a2*, *Sod2*), a T cell-suppressive group (*Cd274*, *Arg2*), and a pro-survival gene (*Cflar*). The gene associations with *Acod1* were then re-analyzed as partial correlations, holding constant either an interferon or an NF-κB score. As shown in Figs 4G–4H, the unadjusted *Acod1* associations with *Slc7a11*, *Arg2*, *Sod2*, *Cflar*, and *Cd274* in MNPs, and with *Slc7a11*, *Sod2*, and *Cd274* in neutrophils, were considered statistically significant, as they exceeded the random background variance generated by shuffling *Acod1* expression. Adjusting for interferon and NF-κB activities partially weakened, but did not nullify, the gene associations in MNPs, while, in neutrophils, they accounted for the *Acod1* associations with *Sod2*.

To validate the above observations, we reanalyzed the DSS colitis dataset, reported by Zhao et al. (22). In that study, mice were administered 3% DSS and cells were collected at day 7. After we independently re-clustered the cells using raw GEO counts, MNPs were split into a freshly recruited monocyte cluster at stage P1 (Ly6C^hi^) and a transitional monocyte cluster at stage P2 (C1q^hi^) (Supplementary Fig. S4A). The same re-clustering led to two neutrophil populations, mostly mature G5a and G5b neutrophils (Supplementary Fig. S4B). As in the case of the Ho et al. dataset (29), *Acod1* expression was elevated in newly arrived Ly6C^hi^ monocytes compared to transitional C1q^hi^ ones and it was almost universally expressed at similar levels by the two neutrophil groups. Moreover, correlation of *Acod1* with the six genes of interest replicated observations made in the Ho et al. dataset (29) for MNPs (Supplementary Fig. S4C) and in part for neutrophils (Supplementary Fig. S4D). Zhao et al. (22), who were the first to link *Acod1* expression with neutrophils in DSS colitis, also published a bulk RNA-seq dataset from an in vitro experiment, in which neutrophils, isolated from the bone marrow of mice, were challenged with LPS in the absence and presence of 4-OI. We used this bulk RNA-seq dataset to determine how itaconate affected the expression of the genes of interest and discovered that 4-OI drove expression of *Slc7a11* and *Slc3a2*, the genes responsible for the cystine/glutamate antiporter System xc^-^, beyond their LPS-induced levels and downregulated the levels of *Sod2* and *Cflar*. The effect of 4-OI on the levels of *Arg2* or *Cd274* was significant but nominal (Supplementary Fig. S4E). As expected, 4-OI supplementation increased genes associated with the NRF2 pathway along with *Acod1*, while downregulating IFN-inducible pro-inflammatory genes (Supplementary Fig. S4F).

Because our observations have linked elevated *Acod1* expression with neutrophil maturity, we explored the association using the publicly available dataset by Xie et al. that established the developmental and mature neutrophil stages (32). To that end, we examined the *Acod1* expression in neutrophil stages, reported by the authors to be present in the bone marrow and blood of healthy as well as *E. coli-*challenged mice. In Supplementary Fig. S5A, *Acod1* expression is negligible in immature stages G1 to G3 but it is switched on as soon as neutrophils transition to mature stage G4. This G3-to-G4 *Acod1* upregulation was independent of the cell compartment and the physiological state of the mice as it was observed in both the bone marrow and blood of healthy and *E. coli*-infected mice (Supplementary Figs. S5B-S5C). The presence of *Acod1* in neutrophils of healthy mice is somewhat surprising as *Acod1* is considered an inflammation-inducible gene; the observation might be suggestive of a special, inflammation-independent *Acod1* function in neutrophil maturation.

Finally, we investigated whether *ACOD1* expression patterns observed in DSS colitis are conserved in the human disease. Subsequently, we analyzed three independent scRNA-seq datasets [TAURUS (33), Kong et al. (34), and Garrido-Trigo et al (23)] involving samples from 90 IBD patients. Because neutrophils are difficult to capture via scRNA-seq, we focused on *ACOD1* presence in the MNP compartment and determined that, despite some patient heterogeneity, *ACOD1* was expressed in the majority of patients and was confined primarily within monocytes (Supplementary Fig. S6A). Because the studies involved different platforms and depth, each dataset was analyzed independently with an identical pipeline and per-cohort effects were combined by random-effects meta-analysis rather than by cell-level integration. With the donor as the unit of inference, *ACOD1* tracked ISG expression in the three datasets (donor-level *ρ*=+0.64, +0.71 and +0.82 for TAURUS, Kong and Garrido-Trigo, respectively). MNPs were clustered according to each dataset authors’ preference and, for each cluster, a monocyte-minus-macrophage identity score and an ISG score were calculated (Supplementary Fig. S6B). Supplementary Fig. S6B demonstrates that *ACOD1* is primarily present in interferon-stimulated, immature monocytes. A within-patient comparison of the transcriptional signatures for *ACOD1*^+^ and *ACOD1*^-^cells further revealed that *ACOD1* characterizes a subset of monocytes with robustly activated pro- and anti-inflammatory programs, alongside upregulated pro-survival pathways that likely allow the cells to survive under severe inflammatory stress (Supplementary Fig. S6C). When *ACOD1* expression was correlated with the six genes of interest (*SLC7A11*, *SLC3A2*, *ARG2*, *SOD2*, *CFLAR*, and *CD274*) and the correlations were then parsed for the presence of interferon or NF-κB, SOD2 and CD274 emerged as a gene signature independently associated with ACOD1 (Supplementary Fig. S6D). Collectively, the above observations suggest that, at the transcriptional level, DSS colitis recapitulated human disease in terms of *ACOD1* expression.

### ERG344 reduces immunosuppression in the TME and suppresses tumor growth in syngeneic cancer models

We first evaluated the therapeutic efficacy of ERG344 in an aggressive form of intraperitoneal carcinomatosis, also used by Weiss et al. to demonstrate that peritoneal tumors elicit Acod1 expression in monocyte and peritoneal macrophage populations (24). In the experiment of Fig. 5A, pretreatment of mice with 1 mpk ERG344 2h prior to B16F10 tumor cell inoculation, reduced the gross tumor burden a week later as evident by the visual inspection of the peritoneal cavity of mice at day 7 post-inoculation. In the images of Supplementary Fig. S7, ERG344 treatment led to a reduced number of disseminated peritoneal tumor nodules. Moreover, ascites collected from vehicle-treated mice exhibited high cellularity, comprising large numbers of tumor and non-tumor cells with >90% viability. In contrast, ascites from ERG344-treated mice contained almost exclusively non-viable tumor cells. Although Acod1 inhibition reduced the tumor burden of mice, it did not prevent eventual mortality. Nevertheless, the median survival increased from 13 days in vehicle-treated controls to 16 days in mice that received ERG344, a statistically significant observation (Fig. 5B).

**Figure 5.**
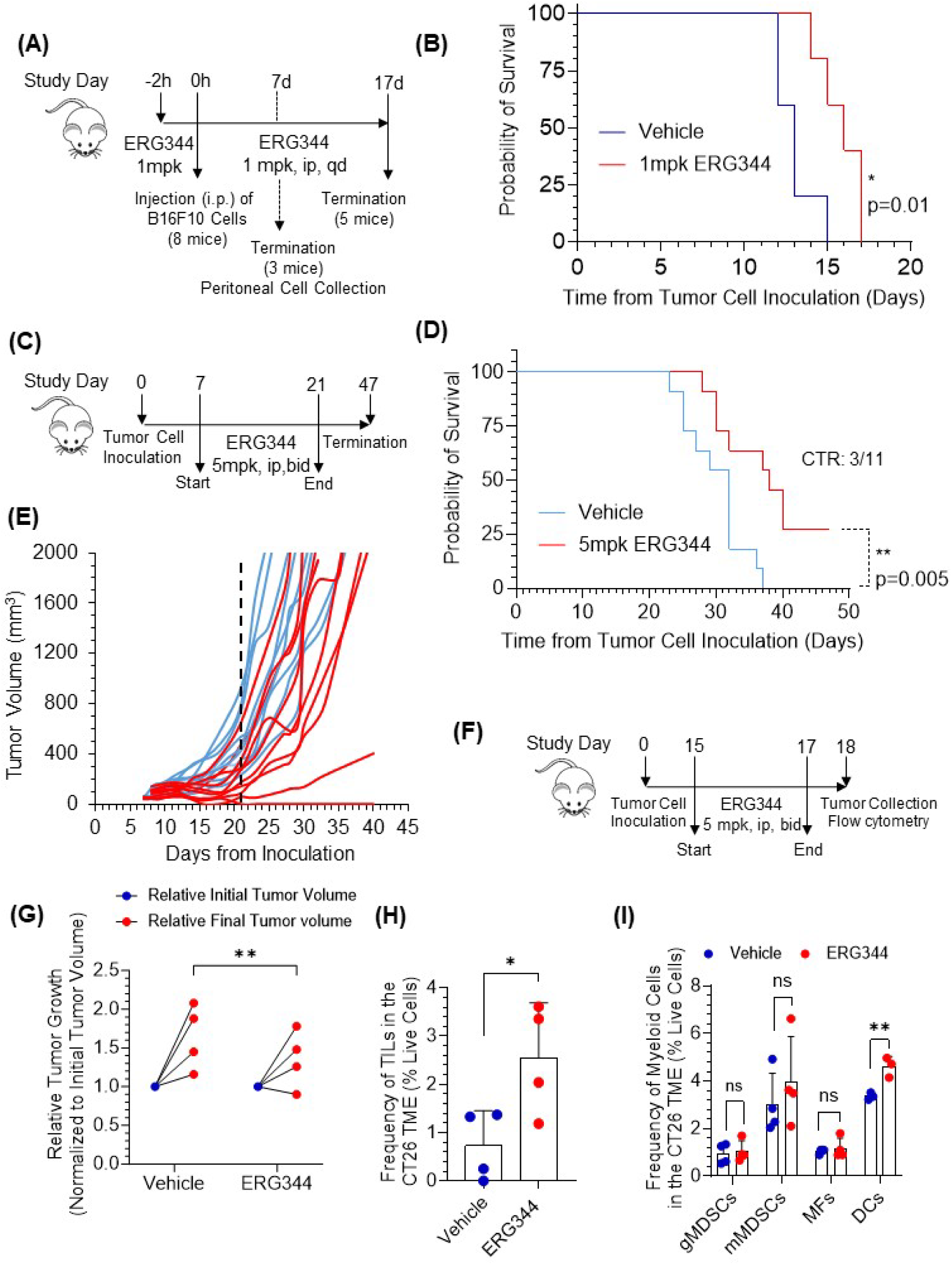
ERG344 suppresses tumor growth and reduces immunosuppression in syngeneic tumor models. (A) Design of the intraperitoneal B16F10 melanoma study (n=8 mice/group). ERG344 administration (1 mpk, ip, qd) was started 2h before tumor cell inoculation. Seven days after inoculation, 3/8 mice were euthanized for peritoneal cell collection. (B) Kaplan-Meier survival curve of B16F10 tumor-bearing mice (n=5 mice/group). (C) Design of the subcutaneous CT26 survival study; two independent experiments; total n=11 mice/group; ERG344 was given ip at 5 mpk, twice a day. (D) Kaplan-Meier survival of CT26 tumor-bearing mice CTR: complete tumor regression. (E) Individual tumor growth curves; dashed line: end of dosing. (F) Design of the short-course CT26 tumor microenvironment (TME) study (n=4 mice/group). ERG344 (5 mpk, ip, bid) on days 15, 16 and 17, tumors profiled on day 18. (G) Change in tumor volume during ERG344 administration in the experiment of (F); tumor volumes were normalized to corresponding ones at the start of treatment. (H) Frequency of tumor-infiltrating lymphocytes (TILs) among live cells in the CT26 TME. (I) Frequencies of granulocytic MDSCs, monocytic MDSCs, macrophages and dendritic cells among live cells in the CT26 TME. Survival curves were analyzed by the log-rank (Mantel–Cox) test. *P<0.05; **P<0.01.

The therapeutic impact of ERG344 was also evaluated in the subcutaneous CT26 colon cancer model (Fig. 5C). In that model, ip administration of 5 mpk ERG344 to mice bearing well-established tumors significantly prolonged overall survival compared to vehicle control mice with the median survival increasing from 31 days in the vehicle-treated animals to 37 days in the ERG344-treated ones (Fig. 5D). More importantly, 27% of the mice (3/11) in the ERG344 group experienced complete tumor regression, although the regression was durable only in 2/11 mice. Individual tumor growth curves, shown in Fig. 5E, continued to diverge well after ERG344 administration was terminated at day 21, indicating a durable effect. To understand how ERG344 treatment affects the tumor microenvironment (TME), mice bearing well-established CT26 tumors were treated briefly with ERG344 (5 mpk on days 15, 16, and 17) and tumors were profiled on day 18 (Fig. 5F). Despite the brevity of administration, ERG344-treated tumors grew slower compared to tumors treated with Vehicle (Fig. 5G). Additionally, flow cytometric analysis of excised tumors revealed a 3.3-fold increase in TILs (Fig. 5H). Detailed myeloid profiling revealed a selective enrichment of mature dendritic cells (CD3^-^CD11b^+^F4/80^-^MHCII^+^CD206^-^), while frequencies of granulocytic myeloid derived suppressor cells (gMDSCs) characterized as CD3^-^CD11b^+^Ly6G^+^Ly6C^lo/int^, monocytic MDSCs (or CD3^-^CD11b^+^Ly6G^-^Ly6C^hi^), and tumor-associated macrophages (CD3^-^CD11b^+^F4/80^+^) remained unchanged (Fig. 5I). The data in Figs. 5H–5I collectively indicate that ACOD1 inhibition converts an immunosuppressive TME into an active, antigen-presenting one that supports antitumor immunity.

### Analysis of *ACOD1* expression in syngeneic cancers

To understand the role of ACOD1 in cancer, we evaluated its gene expression in treatment-naive tumors using a CD45^+^ single-cell atlas comprising 30 tumors across ten syngeneic mouse models (35). Because the publicly available dataset did not include cell-level annotations, we independently derived cell populations from raw count matrices, integrated across the ten models with Harmony to remove tumor-of-origin effects. We determined that MNPs dominated the CD45^+^ populations found in the TME (30%-65% of CD45^+^ cells depending on the model), while neutrophils were not always present and varied greatly (0.1% to 19.4% of CD45^+^ cells) (Supplementary Fig. S8A). In the TME, *Acod1* expression was restricted to the myeloid populations with the *Acod1* gene being detected in 84% of neutrophils and 29% of MNPs. *Acod1* was minimally present in other cell lineages. Following depth-normalization, we determined that *Acod1* expression was highly variable across the MNP compartment of various tumor models peaking at ∼45% of MNPs in CT26 and bottoming out at 1.7% of MNPs in RENCA (Fig. 6A). After applying Leiden clustering with whole transcriptome marker annotation, 120,374 MNPs were resolved into five transcriptional states: an interferon-stimulated monocyte state (Mo_ISG, 22.1%), an MHC class II–high state (MHCII_hi, 20.9%), a pro-tumoral state (Mphi_SPP1, 22.7%), a tissue-resident-like state (Mphi_FOLR2, 26.8%), and a cycling state (Mphi_MKI67, 7.6%) (Fig. 6B). In Fig. 6B, *Acod1* expression was confined primarily within the immature monocyte population characterized by interferon stimulation (Mo_ISG). Following a similar approach, neutrophils were resolved into five states, G5a, G5b, G5c, suppressive, and immature neutrophils (Supplementary Figs. S8B-S8C). Within these states, *Acod1* was ubiquitously present although its mean expression varied with highest value being recorded in state G5b, the state with the highest level of interferon stimulation. Because neutrophils were present in appreciable numbers (>10% of CD45^+^ cells) only in three tumor models (pancreatic Pan02, breast EMT6, and lung LL2) and, in general, neutrophil depletion does not appear to significantly impact tumor growth (15,35), we focused subsequent analyses on the MNP compartment.

**Figure 6.**
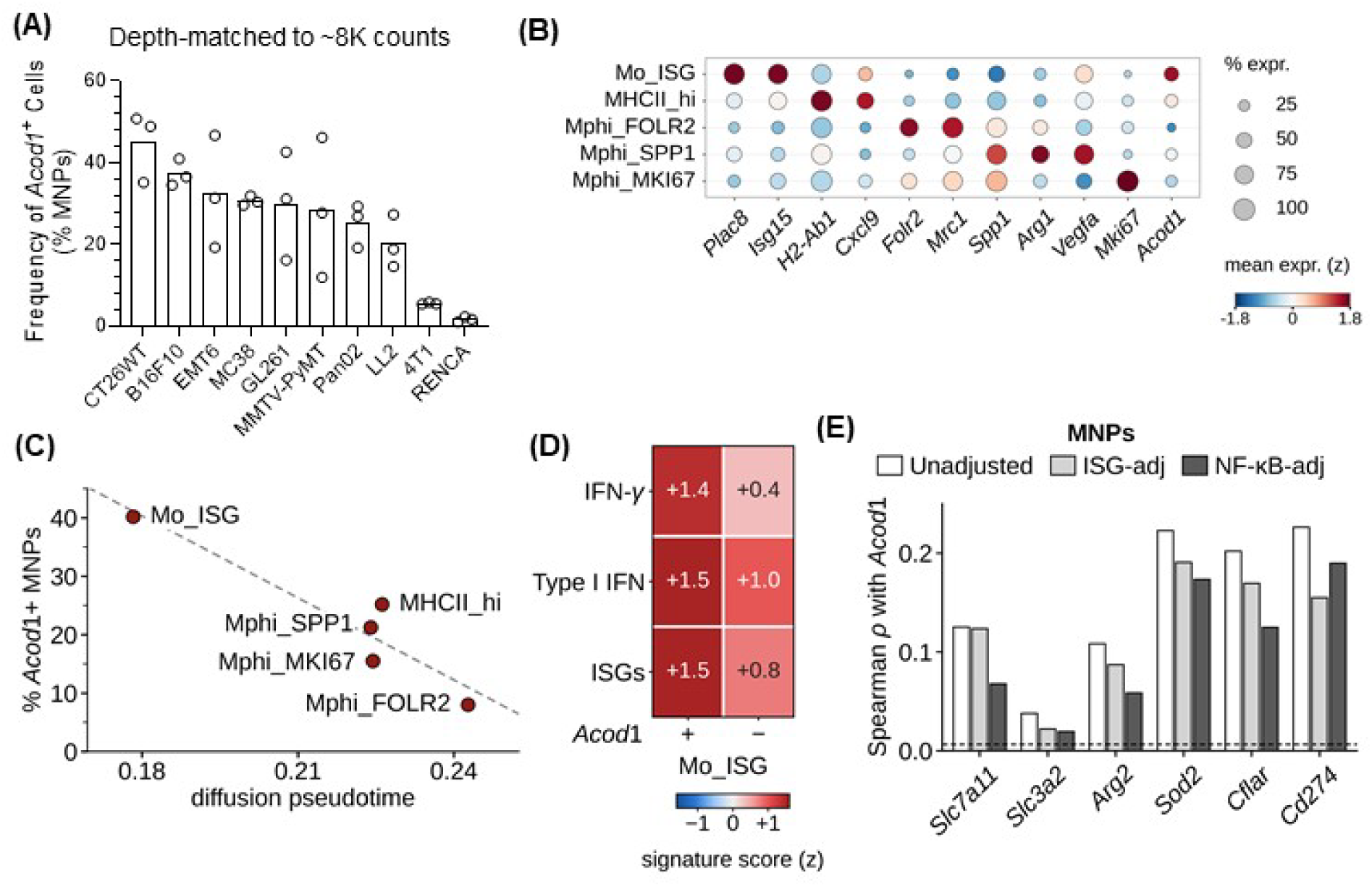
*Acod1* marks interferon-stimulated immature monocytes across ten syngeneic tumor models. Reanalysis of the CD45^+^ single-cell atlas comprising ten syngeneic models and 30 treatment-naive tumors. Cells were re-clustered from raw counts and integrated with Harmony. (A) Frequency of *Acod1^+^* cells among MNPs by model after down-sampling to ∼8,000 counts per cell; each point is one tumor. (B) Dot plot of state-defining markers and *Acod1* across the five MNP states. Dot size: frequency of cells expressing gene. Color: mean z-scored expression. (C) Frequency of *Acod1^+^* MNPs against mean diffusion pseudotime for each MNP state. Dashed line: linear fit for visual purposes only. (D) Interferon signature scores in *Acod1^+^* versus *Acod1^-^* Mo_ISG cells. (E) Spearman correlation of six selected genes with *Acod1* across MNPs, unadjusted and after adjustment for interferon or NF-κB activity. Dashed line, 95th percentile of the permutation null.

The primary objective of our MNP studies was to determine if *Acod1* expression characterized a specific type of monocytes independent of disease state. We first explored what happens to *Acod1* expression with monocyte maturation by building a PAGA graph and calculating diffusion pseudotime (Supplementary Fig. S8D) and then by ordering the five MNP states along a differentiation axis (Fig. 6C). The analysis positioned interferon-stimulated Mo_ISG cells at the start of the developmental trajectory and tissue-resident-like macrophages Mphi_FOLR2 at the end. The remaining three states represented intermediate maturation stages that linked moderately with *Acod1*-rich Mo_ISG in the PAGA graph of Supplementary Fig. S8D. In that differentiation continuum, *Acod1* expression declined along the pseudotime confirming that *Acod1* marks immature monocytes. We then examined the phenotype of *Acod1*^+^ Mo_ISGs. To relate the five MNP states to function we scored each state against five curated gene programs and determined that while MHCII_hi cells were clearly M1-like antigen-presenting macrophages and Mphi_SPP1 cells were pro-tumoral, M2-like macrophages, the Mo_ISG cluster exhibited a mixed gene signature with mildly elevated antigen-presenting, T cell suppressive and angiogenic/hypoxic phenotype (Supplementary Fig. S8E). By applying Spearman rank correlation over the ten models of the study, we observed several trends regarding the frequency of T/NK cells and frequency of various MNP populations and discovered two statistically significant associations between the number of T/NK cells in the TME and MNP clusters: a negative one with proliferating macrophages and a positive one with antigen presenting properties (Mo_ISG+MHCII_hi) (Supplementary Fig. S8F). We also examined the transcriptomic differences between *Acod1^+^* Mo_ISG and *Acod1^-^* Mo_ISG monocytes and determined that the former possessed increased levels of pro- and anti-inflammatory programs as well as survival genes (Supplementary Fig. S8G). When we investigated the type of interferon driving the inflammatory programs of Mo_ISG monocytes, we determined that *Acod1^+^* Mo_ISG cells were influenced by IFN-γ more than *Acod1^-^*Mo_ISG cells suggesting that the former might share a neighborhood with T cells (Fig. 6D). We finally examined the correlation of *Acod1* with the six genes of interest, previously established, and determined that *Acod1* expression was independently associated with all of them with *Sod2* and *Cd274* exhibiting the highest Spearman ρ values (Fig. 6E).

## Discussion

In the last decade, ACOD1 has emerged as a major player in inflammation, while more recently a limited number of reports have assigned a role for the enzyme in tumor immunology (14–16). The studies agree that ACOD1 is an inflammation-inducible enzyme with highly context-dependent properties in pathological settings. Here, we challenge this consensus and, through the use of a novel ACOD1 inhibitor and re-analyses of scRNA-seq human and murine datasets, we demonstrate that ACOD1 is expressed by similar cell populations and operates via a common mechanism in diverse pathological settings such as IBD and cancer.

Despite numerous studies suggesting that ACOD1 has a pathogenic function in DSS colitis, administration of ERG344, an orally bioavailable small-molecule ACOD1 inhibitor, attenuated acute DSS colitis via an on-target effect. To explain the divergence of impact between gene deletion and protein inhibition of ACOD1, we looked at the cell populations that express *Acod1* in colitis in addition to comparing the transcriptomic effects of cell-permeable itaconate to those of ERG344. The picture that emerged did not contradict well-established effects such as the anti-inflammatory role of itaconate in colonic inflammation but rather pointed towards possible ACOD1 functions in colitis progression, which were likely missed in studies utilizing *Acod1* deficient mice. As expected, ERG344-driven inhibition of Acod1 had a proinflammatory effect in colitis as determined by the 2-fold increase in the frequency of pro-inflammatory macrophages within the myeloid population of LP cells. However, the overall impact of Acod1 inhibition in DSS colitis was a beneficial one. We attributed that effect to a dramatic reduction in the number of immature monocytes reaching the site of colonic damage early in the disease, an altered monocyte-to-macrophage maturation later on, and a suppression in the frequency of mature neutrophils, found in the mucosa at the end of the acute testing. These observations, which were obtained through flow cytometry analyses of single cell suspensions of LP from mice treated with vehicle control or ERG344, point to a possible function of ACOD1 in myelopoiesis. Analyses of scRNA-seq datasets confirm that *Acod1* is expressed by neutrophils in the bone marrow of healthy and *E. coli*-challenged mice, where its expression marks the neutrophil transition from an immature (G3) to a mature (G4) stage. It is believed that the G3-to-G4 transition is necessary for the marrow egress of neutrophils. In that light, the ERG344-driven decrease in mature neutrophils points to an Acod1 requirement for maturation of neutrophils and migration to the site of inflammation. Acod1 has recently been implicated in reverse neutrophil migration in which neutrophils leave the site of inflammation and return into circulation (36). However, Acod1 promotes reverse migration and its inhibition would be expected to increase and not suppress the number of mature neutrophils in the mucosa. Integrating the findings of the present study with data reported in the literature suggests that Acod1 exerts both protective and pathogenic effects in colitis. We reason that, while the protective function of Acod1 is the result of itaconate synthesis, the pathogenic properties of the enzyme are linked to temporal production and/or mucosal recruitment of myeloid cells. Mitigating the pathogenic effects of Acod1 via systemic inhibition yields a net therapeutic benefit, while genetic *Acod1* deletion disproportionately disrupts its protective function. Here, it is worth noting a recent study suggesting that, in sepsis, high concentrations of itaconate, reached either through endogenous accumulation or through exogenous administration, exacerbate inflammation and multi-organ failure (37). This observation points to another potential mechanism of action for ERG344.

The association of *ACOD1* with immature monocytes rather than differentiated macrophages has been reported before and confirmed in our re-analyses of scRNA-seq datasets of murine and human colitis (18,23). *ACOD1* expression marks a specific population of newly infiltrating monocytes that are interferon-stimulated and, as a result, have become over-activated and are exhibiting elevated pro- and anti-inflammatory programs. These cells appear stressed with increased levels of survival genes. Notably, identical trends in expression of *Acod1* were seen following re-analysis of a single-cell atlas comprising ten syngeneic tumor models. In those models, *Acod1* expression was correlated with the degree of monocyte maturation with immature monocytes exhibiting the highest level of expression and tissue resident-like macrophages the lowest. This observation was in line with the ERG344 effect on monocyte maturation in colitis and suggestive of an Acod1 function in monocyte production, migration, and maturation under diverse pathological conditions.

Analyses of neutrophils and *Acod1^+^* monocytes revealed common transcriptomic signatures associated with ACOD1 expression independently of interferon- and NF-κB-driven inflammation. The rise in *SOD2* and *CD274* expression along with *ACOD1* held true across most cell types and pathological settings and appeared to be independent of itaconate. While *SOD2* is essential in assisting cells to overcome oxidative stress, *CD274* is an immunosuppressive gene that has been associated with *ACOD1* before (7,18). Notably, our data suggest that, in addition to upregulating CD274, ACOD1 contributes to the rise of an immunosuppressive TME by driving the expression of enzymes such as SLC7A11/SLC3A2 and ARG2, which are known to metabolize cystine and arginine, respectively. It is understood that nutrient-depletion from the TME leads to T cell exhaustion (38,39). In our CT26 colon cancer study, ERG344 administration led to an increase in antigen-presenting DCs and an increase in TILs demonstrating that pharmacological blockade of Acod1 reverses immunosuppression as the transcriptomic signature predicts.

Collectively, our data suggest that ACOD1 functions through a common mechanism in diverse disease settings, which involves myeloid production, differentiation, marrow egress, and/or extravasation. Moreover, ACOD1 expression might drive a specific gene signature in certain myeloid populations, which is responsible for itaconate-independent functions of the enzyme. This cautions against conclusions drawn through itaconate supplementation and suggests that ACOD1 inhibition might be the correct way of interrogating the role of the protein in pathological conditions. Contrary to what has been suggested, we have demonstrated that drugging ACOD1 in inflammatory diseases as well as cancer is a viable therapeutic strategy.

## Materials and Methods

### Materials

Tissue culture media, antibiotics, fetal bovine serum, ultra pure water were from Thermo Fisher. Puromycin dihydrochloride powder was from Sigma Aldrich (Cat. No: P8833). LC/MS grade solvents were from Fisher Scientific. GC/MS-related materials were from Agilent with the exception of methoxyamine hydrochloride (MOX) and N15-leucine, which were purchased from Sigma Aldrich, and MSTFA, which was purchased from Macherey-Nagel. All antibodies for flow cytometry were purchased from Biolegend. Itaconate, used as standard in GC/MS, was from Cayman Chemicals. DSS (MW 36K-50K) was from MP Biomedicals (Cat. 160110). LPS was from Sigma Aldrich (Escherichia Coli 055: B5, Cat. No: L2880).

### Cell lines

Murine colon cancer CT26 cells and murine melanoma B16F10 cells were purchased from ATCC (www.atcc.org) and cultured in Dulbecco’s Modified Eagle’s Medium (DMEM) containing 10% FBS and antibiotics. For in vivo experiments, fresh vials of CT26 and B16-F10 cells were thawed and cultured in 75 cm^2^ T-flasks to 90% confluency prior to tumor cell inoculation. Rat glioma C6 cells were purchased from ATCC and were engineered to overexpress rat ACOD1 by Applied Biological Materials (ABM) Inc. Briefly, cells were transduced at an MOI of 5 for 72h with a lentivirus encoding rat Acod1/Irg1 under a CMV promoter, without a tag or reporter, in a pLenti-GIII-CMV-FireFly Luc-2A-Puro backbone (Cat. No: LVP622083). At 48h post-transduction, cells were placed under puromycin selection at a concentration producing >95% death of wt C6 cells in a prior kill curve. Clonal lines were isolated by drug selection. Transgene overexpression was confirmed by qRT-PCR, by firefly luciferase activity, and functionally by GC-MS quantification of intracellular itaconate. *Acod1^+^* C6 cells were routinely cultured in DMEM containing 10% FBS, antibiotics and 1 μg/mL puromycin.

### Pharmacokinetics

The analysis was performed by Eurofins Panlabs Discovery Services Taiwan, Ltd. Non-fasting pharmacokinetic parameters were obtained after administration of 1 mg/kg i.v. or 5 mg/kg p.o. ERG344 to C57BL/6 male mice (6-8 weeks old). At 3, 10, 30, 60, 120 and 240 min (i.v. dosing) or 10, 30, 60, 120, 240 and 360 min (p.o. dosing), mice were sedated under general inhalant anesthesia (3% isoflurane), and 300-400 µL of blood, obtained by cardiac puncture, were collected in tubes coated with lithium heparin. The plasma was harvested after centrifugation and kept frozen at -80°C until further processing. Each animal was subjected to a single blood draw and 3 animals per time point were used. Plasma was also obtained from 3 naïve (drug-free) mice as blank controls. A calibration curve was generated by spiking drug-free samples with known concentrations of ERG344. Spiked samples and unknown samples were processed together using acetonitrile precipitation and then analyzed by LC-MS/MS. Briefly, proteins present in the plasma were precipitated by acetonitrile, 50 µL of plasma samples were transferred to a 96-well plate, and 150 μL of 0.1 ng/μL of oxybutynin in acetonitrile were also added as internal standard. The samples were analyzed using an AB SCIEX API 3000 (LCMS-010) (ionization mode: electrospray, positive ions) and a Eurospher II 100-5 C18 column (gradient elution with two mobile phases: acetonitrile/formic acid 100/0.2 v/v and water/formic acid 100/0.2 v/v). The column temperature was 30°C, the injection volume was 10 µL, and the run time of the method was 5 min. The reportable linear range of the assay (10-10000 ng/mL ERG344) was determined along with the lower limit of quantification (10 ng/mL ERG344). Plasma concentrations were plotted against time. The fundamental PK parameters of ERG344 after iv or po dosing were determined from the non-compartmental analysis of the data using WinNonlin. The oral bioavailability was also calculated.

### Itaconate Measurement

Itaconate concentrations were estimated in cell lysates of *Acod1^+^*C6 glioma cells, murine sera (24h after the mice were challenged with LPS), and colonic tissue lysates (on day 7 of an acute DSS colitis study). For detection of itaconate in *Acod1^+^* C6 cells, cells were plated onto a 6-well plate or 35 mm tissue culture dishes at a density of 1-2×10^6^ cells/dish in the presence or absence of the ACOD1 inhibitor. Twelve hours later, cells were washed with cold TBS 2X, collected via light scraping, centrifuged, and resuspended in 400 μL ice-cold ultra pure water. Cells were lysed by pulsed sonication (3s, 2x) on ice, 2 μg of N15-leucine were included as internal standard, and 400 μL methanol, previously chilled at -20°C, were added to the cell lysates. The lysates were then allowed to stand on ice for 10 min. Subsequently, 400 μL of dichloromethane, previously chilled at -20°C, were added and the samples were vortexed vigorously and centrifuged at 12,000 g at 4°C for 10 min. The aqueous layers of the cell lysates were collected and dried. Dried materials were dissolved in 20 μL of MOX solution, which was prepared by dissolving 20 mg of methoxyamine HCl (MOX) in 500 μL pyridine for 30 min at room temperature. Metabolites dissolved in MOX were incubated at room temperature for 90 min and subsequently derivatized by adding 80 μL of N-methyl-N-(trimethylsilyl) trifluoroacetamide (MSTFA) at 65°C for 60 min. Approximately 80 μL of derivatized metabolites were transferred into clean, glass GC/MS vials and analyzed by GC-MS using a 7890A GC System (Agilent Technologies) equipped with an Agilent J&W DB-5ms column (30 m, 0.25 mm, 0.25 μm), connected to an 5975C Triple-Axis MS Detector (Agilent Technologies) and outfitted with an autosampler. The MS source was held at 230 °C and the quadrupole at 150 °C. The GC oven temperature was held at 50 °C for 5 min, increased to 240 °C at a rate of 10 °C min^−1^, held at 240 °C for 5 min and increased to 300 °C at a rate of 10 °C min^−1^. A volume of 1 μl was injected in the system in a 1:5 split mode. The amount of itaconate present in the sample was determined after peak integration. A similar methodology was used for determining the amount of itaconate in tissue lysates, which were obtained by mincing longitudinal sections of colonic tissue and then lysing the cells by pulsed sonication in ice-cold ultra-pure water. In the case of assessing itaconate in sera, 5 μL of mouse serum were added into 200 μL of ice-cold ultra-pure water along with 2 μg of internal standard, followed by methanol and dichloromethane as previously described.

### Western Immunoblotting

Approximately 1×10^6^ THP-1 cells were plated onto a 6-well plate and treated with Vehicle or the indicated concentrations of ERG344 for 4h. Cells were collected, washed 2x with PBS, heated in 2X Laemmli sample buffer containing 5% β-ME at 90°C (2 min), and then sonicated (3 pulses, 5s each). Cell lysates were heated for an additional 4 min at 90°C and sonicated again. Cell lysates were resolved by SDS-PAGE and transferred onto nitrocellulose membranes. Membranes were then blocked using 5% non-fat milk in 1X TBST buffer or 3% bovine serum albumin (BSA) in 1X TBST buffer (for detection of phosphorylated proteins) and immunoblotted using appropriate primary and secondary antibodies, diluted in blocking buffer. The probed proteins were detected using SuperSignal West Femto chemiluminescent substrate (Thermo Fisher Scientific). Protein bands were quantified using the software ImageJ. Blots were stripped up to four times using a 200 mM glycine solution (pH 2.0) (1h at room temperature) and Western immunoblotting was repeated.

### RNA isolation, cDNA synthesis and qRT-PCR

Total RNA was isolated from mouse colon using the MagMAX mirVana Total RNA Isolation Kit (Thermo Fisher, A27828) on a MagMAX Express-96 magnetic particle processor, including on-instrument DNase treatment, according to the manufacturer’s protocol. RNA was normalized with nuclease-free water to a maximum of 4 µg in 20 µl and reverse-transcribed using the High Capacity cDNA Reverse Transcription Kit (Applied Biosystems, 4368813) in a 40 µl reaction (25 °C for 5 min, 37 °C for 120 min, 85 °C for 5 min, hold at 4 °C). Quantitative PCR was performed in technical duplicate in MicroAmp Optical 96-well plates containing 9 µl TaqMan Fast Advanced Master Mix (Applied Biosystems, 4444557) with the respective probe and 1 µl cDNA per well, on a QuantStudio 6 Flex system (Applied Biosystems). Cycling conditions were 95 °C for 20 s, followed by 40 cycles of 95 °C for 1 s and 60 °C for 20 s. Inventoried TaqMan Gene Expression Assays were used for *Actb* (Mm02619580_g1), *Il1a* (Mm00439620_m1), *Il1b* (Mm00434228_m1), *Il6* (Mm00446190_m1), *Tnf* (Mm00443258_m1), *Isg15* (Mm01705338_s1), *Acod1*/*Irg1* (Mm01224532_m1) and *Hmox1* (Mm00516005_m1). qRT-PCR assays were performed by DC3 Therapeutics (South San Francisco, CA). Relative expression was calculated by the comparative C_T_ [2^(−ΔΔC^_T)_] method with *Actb* as the reference gene.

### Endotoxemia In Vivo Model

Female C57BL/6 mice (ENVIGO, 6-8 weeks old, Strain: 3928) were randomized into treatment groups (n=4 mice/group) based on average body weight using GraphPad Prism. The mice were pretreated with 50 mpk ERG344 for 30 min prior to challenging the mice with 1.5 mpk LPS; they were treated again with ERG344 eight hours after the LPS challenge. Sera were collected 2h after LPS injection and at the end of the experiment. The experiment was performed by Washington Biotechnology Inc. Itaconate was assessed at the terminal sera, while sera collected at 2h-post LPS injection were used to assess TNFα in circulation. Serum TNFα was estimated from 10 μL serum using an ELISA kit from Biolegend (ELISA Max Deluxe Set Mouse TNFα, Cat. No: 403904).

### Acute DSS-Induced Colitis Model

Female C57BL/6 mice (Charles River Laboratories, 18-20 g each, 8-9 weeks old, Strain: #4338) were acclimated for 7 days ear-tagged, and randomized into treatment groups (n=5/group) based on average body weight using GraphPad Prism. Colitis was induced with 2% (w/v) DSS in drinking water ad libitum. Treatment was initiated 24h after first exposure to DSS. Mice were treated with either ERG344 or with Vehicle (PBS) control for the indicated number of days. They were observed daily for clinical signs of colitis and euthanized 24h after the last treatment. For tissue collection, colons were excised at the ileocecal junction, measured for length, and opened longitudinally to clear fecal content before weighing. Baseline measurements were obtained from naive control mice (n=3 mice) at the beginning of the work and then treated as historical values in subsequent studies. Tissues were stored in tissue storage solution (Miltenyi Biotec) until they were processed for flow cytometry, typically within two hours from collection. For all studies, protocols were prepared by the CRO (Washington Biotechnology Inc.). The mice were scored for signs of colitis according to the following parameters: Body weight loss (0: ≤1%, 1: ≤5%, 2: ≤10%, 3: ≤15%, 4: >15%); Stool consistency (0: separate hard lumps, 1: sausage-like, 2: soft blobs, 3: watery); Anal prolapse (0: not visible outside the body, 1: visible outside the body); Fecal blood (0: no blood in stool; 1: blood in stool). Disease Activity Index was calculated as the sum of all scores. DSS colitis was also studied in female Acod1^-/-^ mice from Jackson Laboratories (C57BL/6NJ-Acod1em1(IMPC)J/J, Strain: #029340,) using the previously described protocol. The experiments were performed by Washington Biotechnology Inc.

### Syngeneic Tumor Model Studies

<u>(A) *Intraperitoneal B16F10 Carcinomatosis Model*</u>. Female C57BL/6 mice (Charles River Laboratories, 8-9 week-old, Strain: #4338) were randomized based on body weight into two groups (n=8/group) receiving either vehicle control or 1 mpk ERG344. Treatment started with a single dose two hours prior to injecting the mice i.p. with 1×10^6^ B16F10 melanoma tumor cells in PBS. The treatment continued with daily, single dose treatments until death. Seven days after the tumor injection, 3/8 mice were euthanized, ascites were withdrawn, and the peritoneal cavities were inspected for evidence of tumor nodules. <u>(B)</u> <u>CT26 colon cancer model.</u> Female BALB/c mice (Charles River Laboratories, 6-8 week-old, Strain: #028) were injected subcutaneously with 1×10^5^ CT26 tumor cells in PBS containing 20% Matrigel in the right flank and tumors were allowed to grow over a period of 7-14 days to a predetermined initial tumor volume. For the survival study, the tumors were grown to an average of 100-120 mm^3^, whereas, for the TME profiling study, the tumors were grown to an average of 300 mm^3^ before they were randomized using the statistical software GraphPad. ERG344 was administered i.p., twice a day (bid) at 5 mpk with the treatment starting one day after animal randomization. Mice in the vehicle control group received PBS. Mice were observed daily for clinical signs of stress. Tumor volumes and body weights were assessed every other day. Mice were euthanized when tumor volumes were ≥2000 mm^3^, as measured using a digital caliper and the following formula: ½(Length x Width^2^). Additional criteria for euthanasia included signs of distress, extensive tumor laceration and weight loss of >20% of body weight. The experiments were performed by Washington Biotechnology Inc.

### Flow Cytometry Assays

Single cell suspensions were obtained from excised, minced colonic tissues or tumors using a Lamina Propria Dissociation Kit (Miltenyi, Cat. No: 130-097-410) for the former or a Tumor Dissociation Kit (Miltenyi, Cat. No: 130-096-730) for the latter according to the manufacturer’s instructions. The cells were subsequently collected, washed with PBS, and stained with Zombie Violet fluorescent dye (Biolegend) per manufacturer’s instructions for live/dead discrimination. Live cells stain negative for Zombie. After removal of the dye, the cells were re-suspended in flow cytometry staining buffer (PBS containing 4% FBS and 2 mM EDTA) and blocked with TruStain FcX (mouse or human, Biolegend) for 5 min at 4°C. The cells were subsequently exposed to a cocktail of antibodies against extracellular biomarkers (if any) for 30 min at 4°C, washed with staining buffer 2X, and fixed using 2% paraformaldehyde in PBS (10 min at 4°C). At the end of the fixation period, the washing step was repeated. If the experiment included intracellular biomarkers, the cells were permeabilized using a permeabilization buffer (Invitrogen), stained with a cocktail of antibodies against the intracellular biomarkers (30 min at 4°C), washed with the permeabilization buffer 2X and resuspended in staining buffer. Biomarker analysis was performed using a BD FACSCanto II flow cytometer. The experiment typically included appropriate isotype control sample(s). Data were analyzed using FlowJo v10. The following general <u>Gating Strategy</u> was employed: The cells were sequentially gated for size using forward versus side scatter (FSC vs SSC), for singularity (only single cells were included), and live cells (Zombie negative cells were selected). Isotype control staining of a biomarker was then used as a threshold to determine its expression in the immune cells of interest.

### Bioinformatics

Computational reanalysis (murine colitis) was performed on four publicly deposited Gene Expression Omnibus (GEO) datasets: GSE148794 (primary scRNA-seq, murine DSS colitis), GSE312526 (independent scRNA-seq DSS confirmation), GSE137539 (scRNA-seq neutrophil maturation reference), and GSE312519 (bulk RNA-seq of bone-marrow neutrophils treated with LPS ± 4-octyl itaconate). *Acod1* (often annotated as *Irg1*) was used strictly as a transcriptional marker rather than a quantitative measure of itaconate. Because samples were heavily pooled, inferences are pseudoreplicated at the cellular level. *Single-Cell Processing and Annotation*. Standard quality control filters (minimum gene counts, maximum UMI limits, and mitochondrial read thresholds) were applied. Counts were normalized (10,000 per cell), natural-log-transformed, and clustered using the Leiden algorithm. Granulocyte and mononuclear phagocyte (MNP) compartments were isolated and sub-clustered independently. Annotations were derived from source publication markers and the Kang et al. framework (30). Crucially, *Acod1* was excluded from all module scores and signatures used to define cell types or clusters. *Differential Expression and Acod1 Classification*. In the MNP compartment, cells with ≥1 raw *Acod1* UMI were classified as *Acod1*^+^; in neutrophils, *Acod1* was modeled as a continuous gradient. Per-gene contrasts were evaluated using two-sided Fisher’s exact tests for detection rates and log₂ fold-changes (with bootstrap standard errors) for mean expression. To control for general cellular activation, contrasts were also performed exclusively within the upper quartile of an interferon-stimulated gene (ISG) score. Stringent limits-of-detection safeguards were applied to exclude spurious artifacts (e.g., *Arg1* in neutrophils). *Regression Models and Partial Correlations.* Transcriptome-wide *Acod1* associations were estimated using Poisson rate models fitted to raw counts, incorporating log total counts as an offset to structurally control for sequencing depth. To distinguish *Acod1*-specific networks from generic inflammation, expression was additionally modeled against an NF-κB module score and negative-control targets; genes were only retained if their *Acod1* coefficient survived this adjustment. Rank-based partial Spearman correlations were used for per-gene analyses. *Permutation Calibration*. Because shared sequencing depth alone can induce positive correlations in sparse data, associations were not tested against a null of zero. Instead, an empirical null was generated by randomly permuting *Acod1* expression within library plates (500 permutations) with library plate defined as one of the 60 separately indexed sequencing libraries in GSE148794 (10 per time point), each prepared from single cells FACS-sorted into 384-well plates. Only associations exceeding the 95th percentile of this within-plate null were considered statistically significant. *Cross-Dataset Stage Assignment*. Monocyte maturation staging was anchored to GSE148794 centroids using smoothed module scores. Conversely, machine-learning classifiers for neutrophil maturation failed to transfer reliably across datasets due to transcriptomic extrapolation limits. Consequently, neutrophil stage analyses relied entirely on direct, classifier-independent marker measurements (e.g., absence of proliferation markers and elevated ISGs) rather than imputed stage labels*. Bulk RNA-seq Analysis*. Differential expression for GSE312519 was performed using pydeseq2. To strictly isolate the transcriptional effect of 4-octyl itaconate (4-OI) against a background of acute inflammation, the primary analytical contrast was restricted to the LPS + 4-OI arm against the LPS-only arm. *Robustness and Reproducibility*. Primary estimates were subjected to a rigorous battery of sensitivity checks, including binomial depth matching, alternative distribution models, and leave-one-out refits. Analyses were performed in Python (Scanpy). Fully reproducible computational notebooks, environment specifications, and per-cell metadata will be deposited publicly at a later date.

### Statistics

Unless otherwise indicated, experimental data are presented as mean±SD, and group comparisons were made using an unpaired, two-tailed Student’s *t*-test. For the DAI and body-weight time courses, vehicle- and ERG344-treated groups were compared at each timepoint. The qRT-PCR comparisons were corrected across genes by the Benjamini–Hochberg procedure and are reported as FDR *q* values. Survival was compared by the log-rank (Mantel– Cox) test. Significance is indicated as *P<0.05, **P<0.01, ***P<0.001, ****P<0.0001; ns, not significant. Statistical methods for the computational re-analyses are given under *Bioinformatics*.

## Supporting information

All Supplemental Figures

## Acknowledgment

A.E.P. acknowledges financial support from the Vitreous State Laboratory at The Catholic University of America (internal funding).

## Data and code availability

The study analyzed only publicly available data. Murine datasets were obtained from the Gene Expression Omnibus under accessions GSE148794 (Ho et al., longitudinal DSS colitis, Ref. 29), GSE312526 (Zhao et al., DSS colitis scRNA-seq, Ref. 22), GSE312519 (Zhao et al., bulk RNA-seq of bone-marrow neutrophils, Ref. 22), GSE137539 (Xie et al, neutrophil developmental atlas, Ref 33), and GSE307143 (Wang et al., the syngeneic CD45^+^ atlas, Ref. 36). Human inflammatory bowel disease datasets were obtained from GSE282122 (TAURUS of Ref. 34; also, processed, annotated data (.h5ad) and the sample-pairing list can be found at Zenodo with doi:10.5281/zenodo.13768607), GSE214695 (Garrido-Trigo et al, Ref. 23), and SCP2959 (Single Cell Portal), phs003943.v1.p1 (dbGaP) of the Broad Institute of MIT and Harvard (Kong et al., Ref. 35). Analysis notebooks, environment specifications and per-cell metadata supporting the computational analyses will be deposited at GitHub. They are currently available from the corresponding author.

## Ethics

Animal model studies were conducted at Washington Biotechnology, Inc., a contract research organization (CRO) located in Baltimore, MD. The studies were approved by the local Institutional Animal Care and Use Committee (IACUC). The animal research facility was registered with the US Department of Agriculture, had an Animal Assurance Statement with the Office of Laboratory Animal Welfare (OLAW), and was AAALAC accredited. Pharmacokinetics studies were executed by Eurofins Panlabs Discovery Services Taiwan, Ltd. The work was performed in accordance with the Guide for the Care and Use of Laboratory Animals (National Academies Press, Washington, DC) using an AAALAC accredited facility. The study protocol was approved by the Eurofins Panlabs Discovery Services Taiwan, Ltd IACUC.

