## Supplementary material for "Small-molecule ACOD1 inhibition attenuates colitis and tumor growth through a shared myeloid mechanism, diverging from genetic deletion": All Supplemental Figures

### Supplementary Figure Legends

**Figure S1. ERG344 Properties.** (A) Reaction catalyzed by ACOD1: decarboxylation of *cis*-aconitic acid to itaconic acid. (B) Schematic of the inflammation-associated break in the TCA cycle that diverts *cis*-aconitate to itaconate. (C) Pharmacokinetic parameters of ERG344 in C57BL/6 mice after a single intravenous dose (1 mpk, upper table) or oral dose (5 mpk, lower table). $t_{1/2}$, elimination half-life; $C_0$, extrapolated initial concentration; $C_{max}$, peak concentration; $T_{max}$, time to peak; $AUC_{last}$ and $AUC_{inf}$, area under the curve to the last measurable timepoint and extrapolated to infinity; $AUC_{extr}$, percentage of $AUC_{inf}$ obtained by extrapolation; $AUC/D$, dose-normalised AUC; MRT, mean residence time; $V_{ss}$, volume of distribution at steady state; CL, clearance; F, oral bioavailability. (D) Serum TNFα 2 h after LPS challenge in mice given vehicle or ERG344 (50 mpk, i.p.). Bars are mean±SD; each circle is one animal (n=4 mice/group). ns, not significant.

**Figure S2. Effects of ERG344 on colon morphology and mucosal myeloid populations during DSS colitis.** (A, B) Resected colons on day 7 of DSS from (A) wt C57BL/6 and (B) *Acod1*<sup>-/-</sup> mice treated with vehicle or ERG344 (5 mpk, i.p.). Each image is one animal (n=5 mice/group). The corresponding studies are found in Fig. 2. (C,D) Frequency of (C) transitioning (CD11b<sup>+</sup>CD11c<sup>-</sup>CX3CR1<sup>int</sup>CD64<sup>-</sup>MHCII<sup>+</sup>) monocytes and (D) mature resident (CD11b<sup>+</sup>CD11c<sup>-</sup>CX3CR1<sup>hi</sup>CD64<sup>+</sup>) macrophages among lamina propria (LP) CD11b<sup>+</sup>CD11c<sup>-</sup> cells on day 4 of DSS; study design in Fig. 3A. (E) Relative expression of inflammatory genes and *Hmox1* in colonic tissues on day 4, normalized to *Actb*. (F) Colon mass-to-length ratio on day 4 in untreated mice and in DSS-challenged mice given vehicle or ERG344. Ratios from DSS-naïve animals are historic values shown for reference. In (C–F), bars are mean±SD; each circle represents one animal (n=4 mice/group). ns, not significant; \*\*P < 0.01.

**Figure S3. Transcriptional characterization of *Acod1*<sup>+</sup> mononuclear phagocytes (MNPs) in murine DSS colitis.** Reanalysis of the longitudinal DSS colitis dataset of Ho et al. (29) (A) Expression of annotating genes across the six re-clustered MNP states. Dot size: percentage of cells expressing; color: scaled mean expression. (B) Within-timepoint Spearman correlation between *Acod1* expression and the interferon-stimulated gene (ISG) score across the disease time course. (C) Expression of genes grouped by functional program in *Acod1*<sup>+</sup> versus *Acod1*<sup>-</sup> monocytes (Ly6C<sup>hi</sup>, ISG-Mono, and transitional MoDM clusters combined). Dot size: percentage expressing; color: mean log<sub>10</sub>(CP10K). \*q < 0.05; †not detected. (D) Fold change in expression of nine inflammatory genes per 1-SD increase in the interferon score (left) or NF-κB score (right), with 95% confidence intervals. Filled circles, q < 0.05.

**Figure S4. Validation of transcriptional findings in an independent colitis dataset.** (A, B) Reanalysis of the DSS colitis dataset of Zhao et al. (3% DSS, day 7) (22). Expression of marker genes and *Acod1* across (A) the two mononuclear phagocyte (MNP) clusters comprising freshly recruited Ly6C<sup>hi</sup> monocytes (P1) and transitional C1q<sup>hi</sup> monocytes (P2) and (B) the two neutrophil states (G5a and G5b). Dot size: percentage expressing; color: mean expression. (C, D) Spearman correlation of the six selected genes with *Acod1* in (C) MNPs and (D) neutrophils, shown unadjusted and after adjustment for interferon (ISG-adj) or NF-κB (NF-κB-adj) activity. Dashed line, 95th percentile of the permutation null. (E) Normalized counts for the six genes in bone marrow neutrophils cultured with vehicle, LPS, or LPS plus 4-octyl itaconate (4-OI), from the bulk RNA-seq dataset of Zhao et al. (22). \*P < 0.05; \*\*P < 0.01; \*\*\*P < 0.001; \*\*\*\*P < 0.0001. (F) Heatmap of NRF2/antioxidant, IFN-GTPase and NF-κB/cytokine target genes across LPS and LPS+4-OI replicates, shown as row z-scores from the previously described bulk RNA-seq dataset.

**Figure S5. *Acod1* is switched on at the immature-to-mature transition during neutrophil differentiation.** Reanalysis of the neutrophil developmental atlas of Xie et al. (32) (A) Expression of maturation-stage markers and *Acod1* across neutrophil stages G1–G5c in bone marrow and blood of healthy and *E. coli*-infected mice. Data are depth-matched to 2,000 UMI per cell; cell numbers per group are given at right. Dot size: percentage expressing; color: mean expression scaled within each gene. Dashed box highlights *Acod1*. (B, C) Percentage of *Acod1*<sup>+</sup> cells at each neutrophil stage in (B) healthy and (C) *E. coli*-infected mice. Bars are mean±SD. Dashed boxes highlight the G3-to-G4 transition. Light blue, bone marrow; dark blue, blood.

**Figure S6. ACOD1 marks interferon-stimulated immature monocytes in human IBD.** Reanalysis of three independent scRNA-seq cohorts [TAURUS (33), Kong et al. (34), Garrido-Trigo et al. (23)] comprising 90 patients. (A) Number of donors per cohort by monocyte and ACOD1 status. (B) Interferon-stimulated gene score plotted against the monocyte-minus-macrophage identity score for each mononuclear phagocyte (MNP) cluster, with clusters annotated as defined by each dataset's original authors. Point size: percentage of ACOD1<sup>+</sup> cells; color: cohort. (C) Expression of genes grouped by functional program in ACOD1<sup>+</sup> versus ACOD1<sup>-</sup> cells, compared within patients. Dot size: percentage expressing; color: mean log1p(CP10K). Black asterisk, $q < 0.05$ by paired Wilcoxon test ($n=24$ patients); blue asterisk, lower in ACOD1<sup>+</sup> cells. (D) Spearman correlation of the six selected genes with ACOD1 in monocytes, unadjusted and after adjustment for interferon or NF-κB activity. Dashed line, 95th percentile of the permutation null.

**Figure S7. Gross peritoneal tumor burden in the intraperitoneal B16F10 model.** Peritoneal cavities of mice 7 days after intraperitoneal inoculation of B16F10 melanoma cells, treated with vehicle (upper panel) or ERG344 (1 mpk, i.p., daily; lower panel). Each image is one animal ($n=3$ mice/group). Red arrows indicate macroscopic tumor nodules.

**Figure S8. Composition and transcriptional states of myeloid cells across ten syngeneic tumor models.** Reanalysis of the CD45<sup>+</sup> single-cell atlas of Wang et al. (30 tumors, ten models) (35). (A) Frequency of mononuclear phagocytes (MNPs) and neutrophils as a percentage of CD45<sup>+</sup> cells in each model. (B) Distribution of neutrophils across the five re-clustered neutrophil states. (C) Expression of annotating genes and *Acod1* across the neutrophil states. Dot size: percentage expressing; color: mean z-scored expression; \*states defined in this study. (D) PAGA connectivity graph of the five MNP states (left, node area proportional to cell number, root set to Mo\_ISG) and the corresponding connectivity matrix (right). Positions along the x-axis give mean diffusion pseudotime. Based on 120,374 MNP cells from ten models, integrated with Harmony across 30 samples. (E) Scores for five curated functional programs across the five MNP states, z-scored across states (96,302 depth-matched cells). (F) Spearman correlation between the frequency of each MNP state and the frequency of T/NK cells across the ten models. (G) Expression of genes grouped by functional program in *Acod1*<sup>+</sup> versus *Acod1*<sup>-</sup> Mo\_ISG cells (9,974 versus 11,304 cells). \*$q < 0.05$.

**Figure S1**

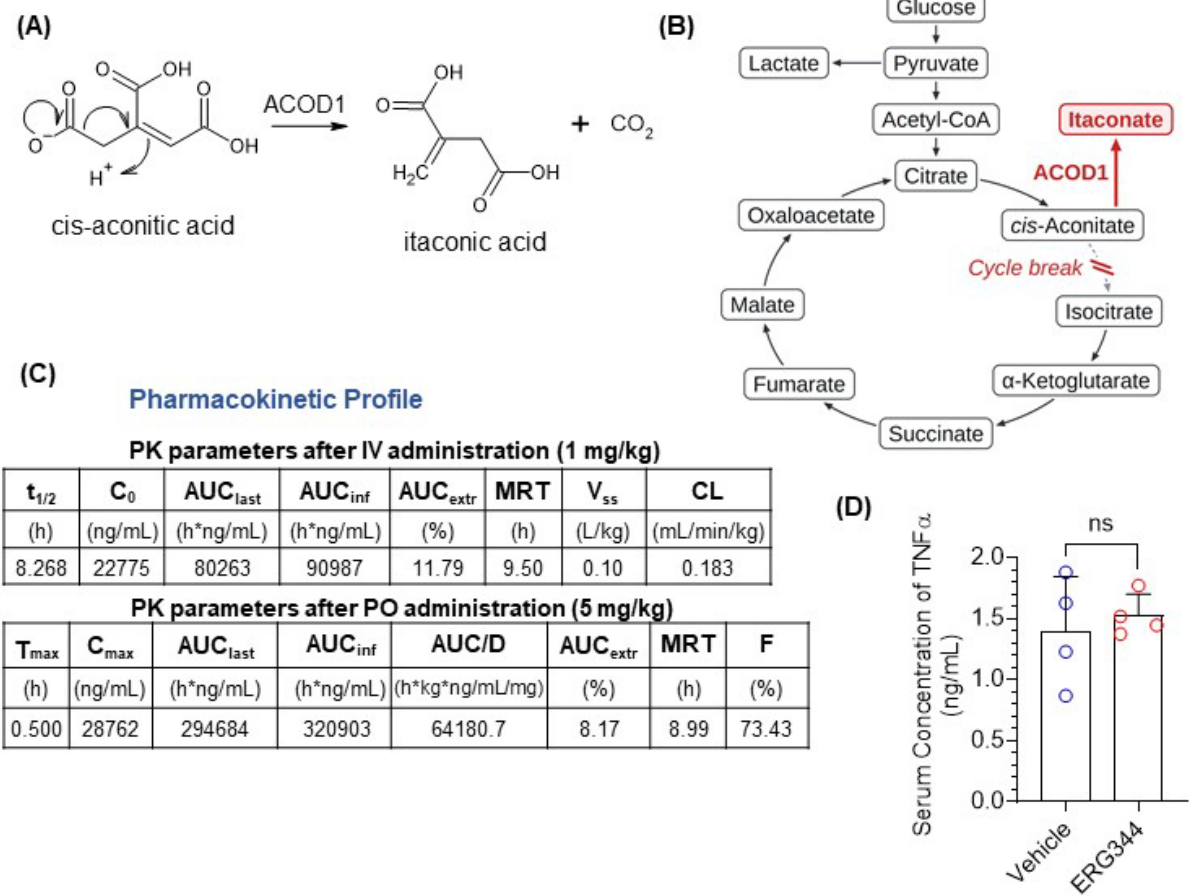

**Figure S2**

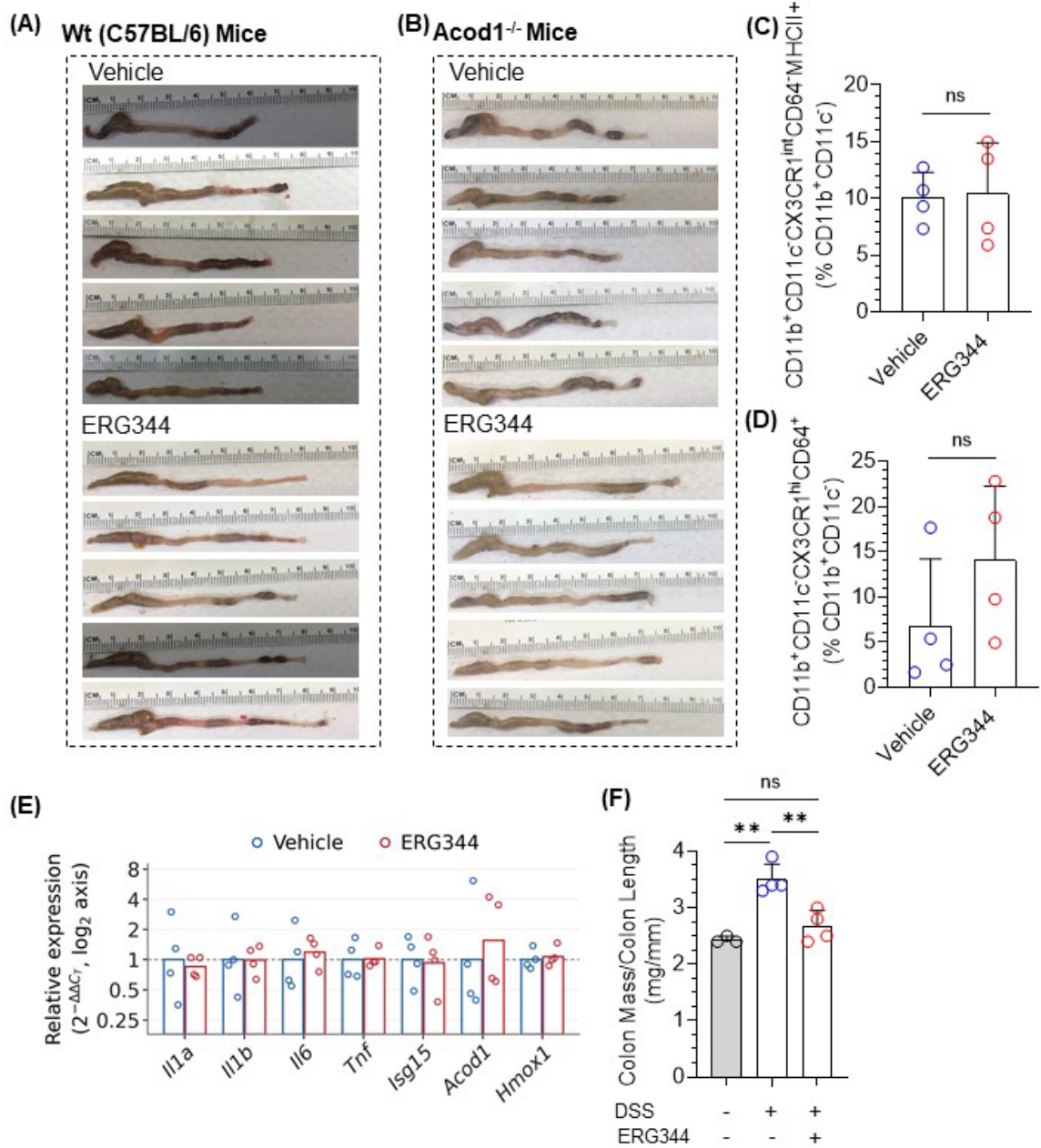

Figure S3

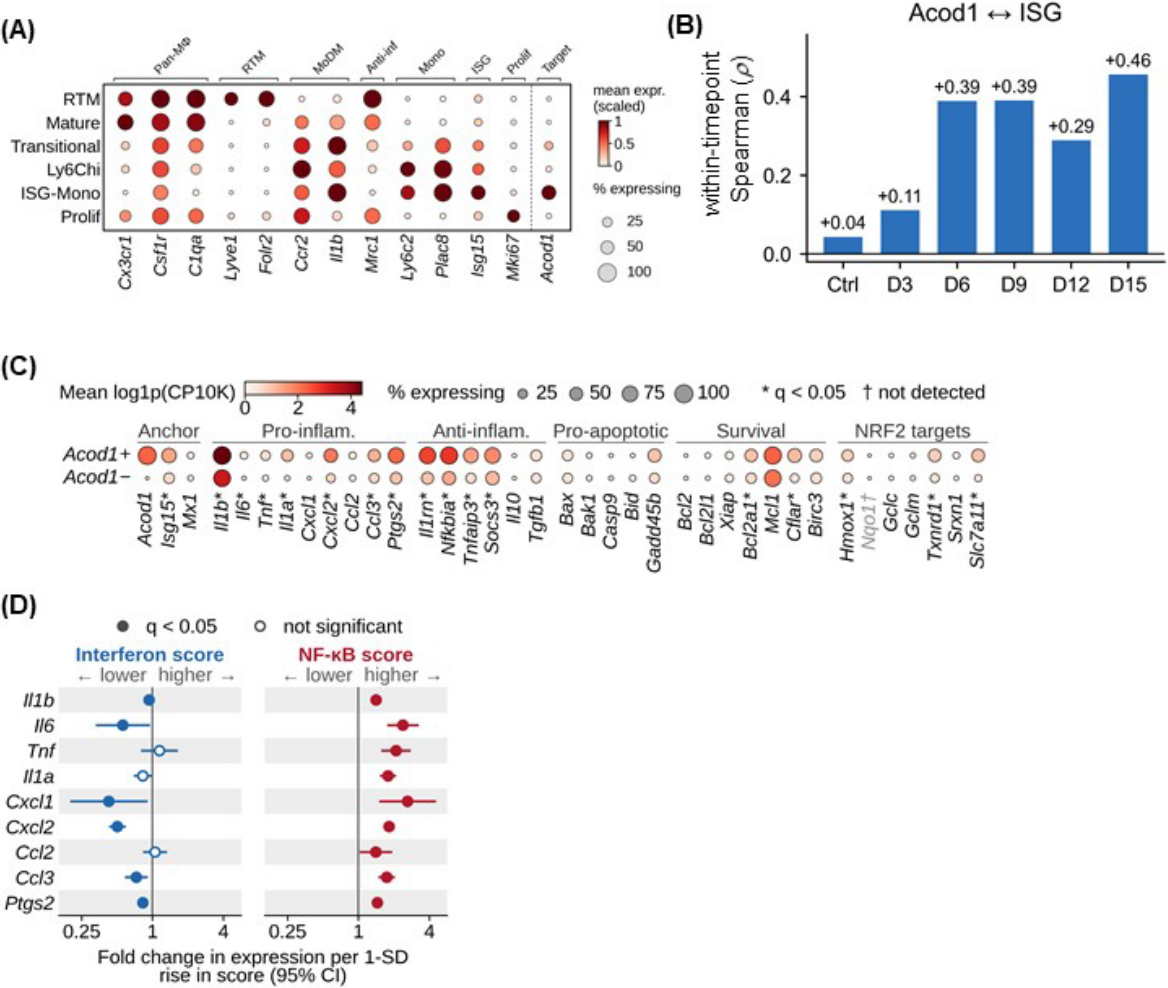

**Figure S4**

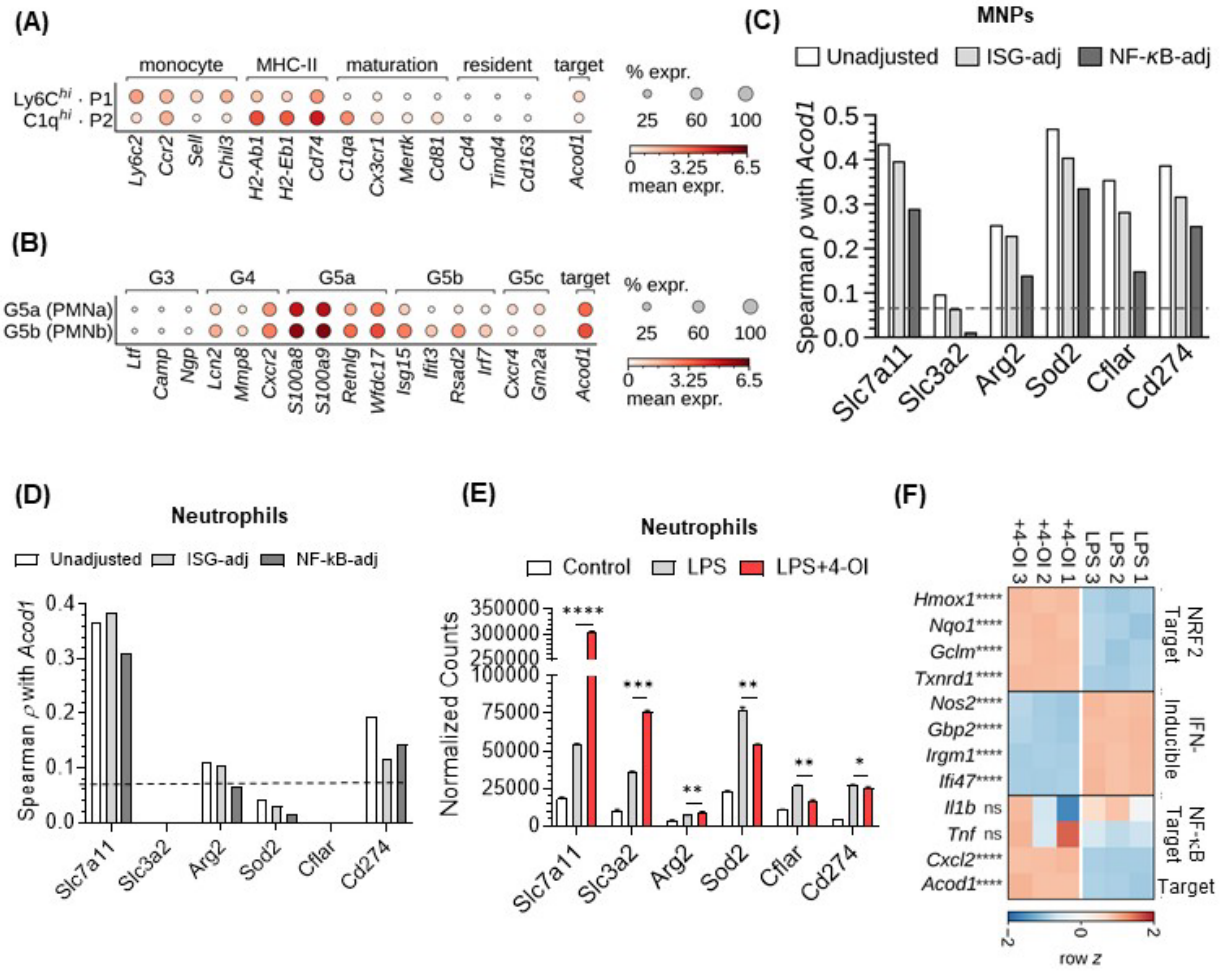

Figure S5

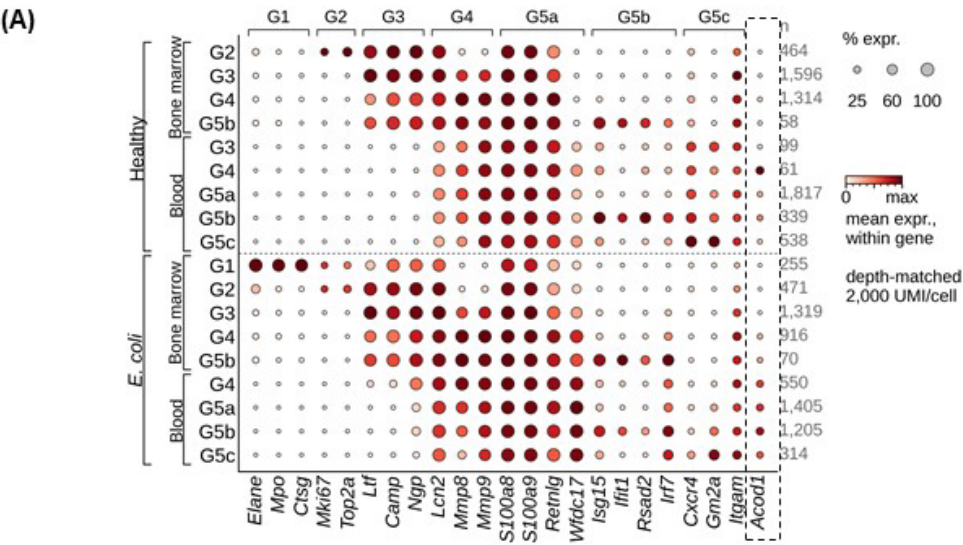

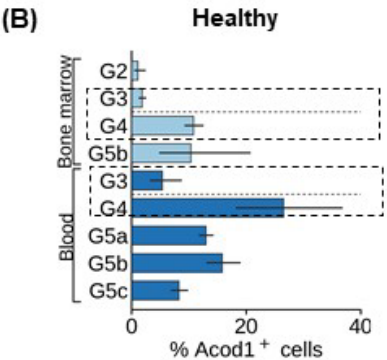

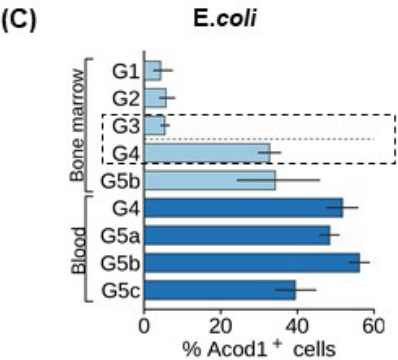

Figure S6

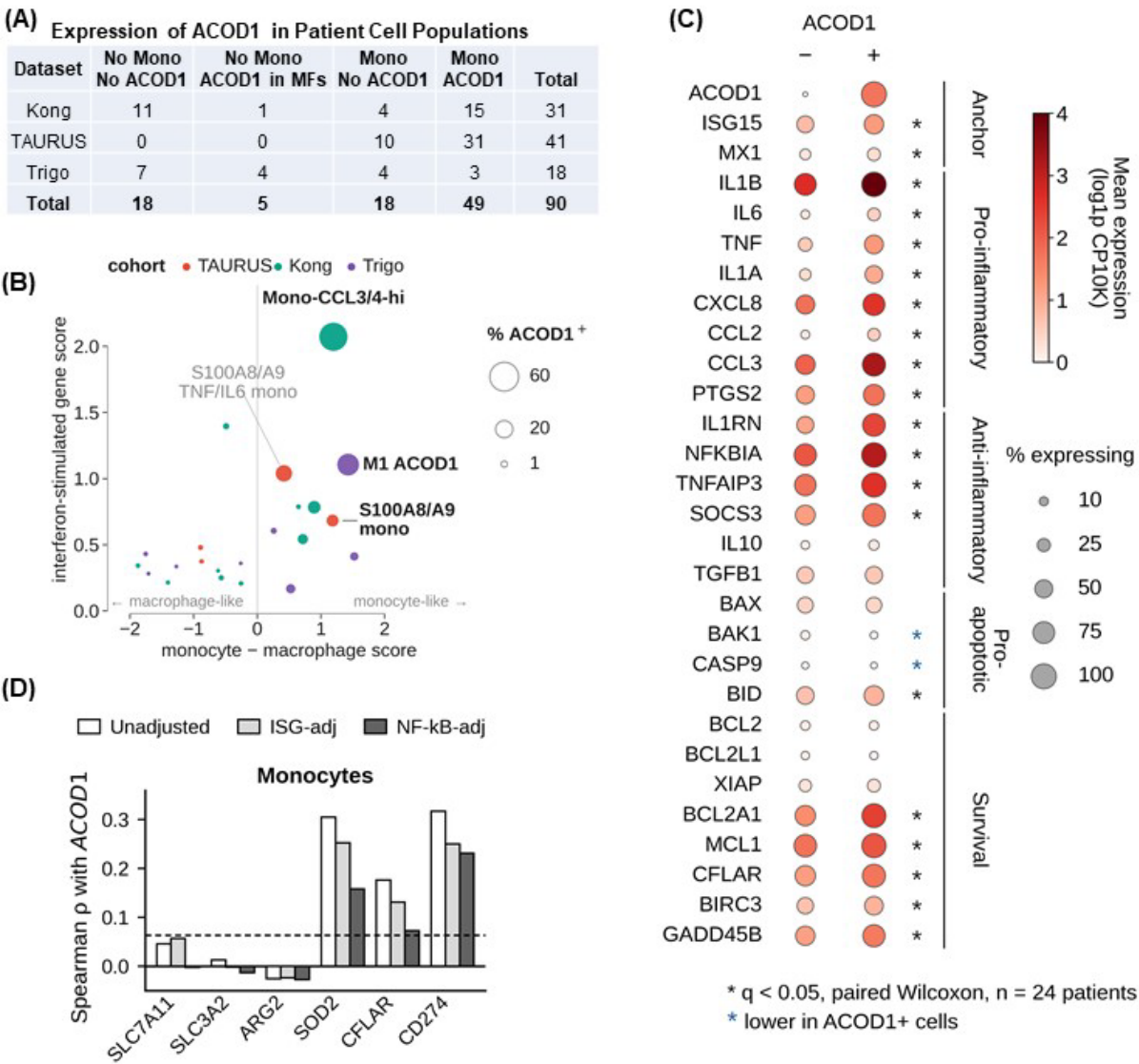

**Figure S7**

**Vehicle**

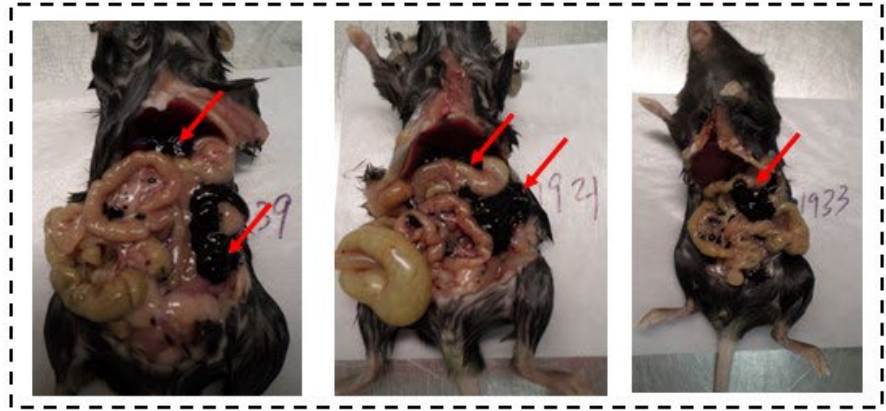

**ERG344**

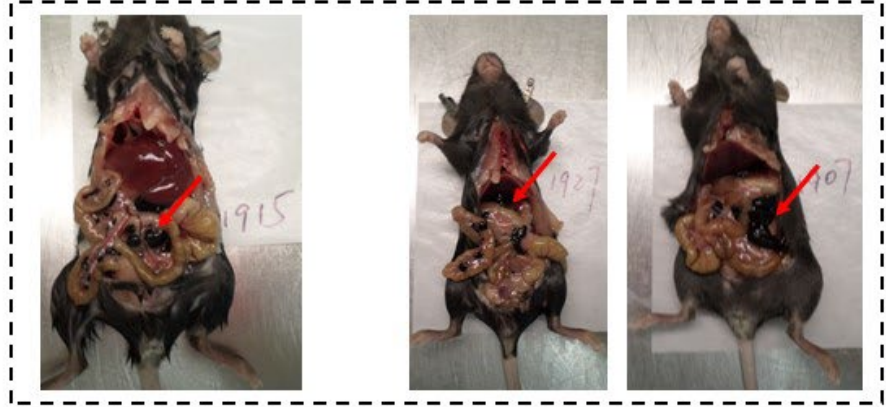

Figure S8

(A)

| Cluster/Tumor | Frequency of Cell Population (% CD45+ Cells in the TME) |  |  |  |  |  |  |  |  |  |
| --- | --- | --- | --- | --- | --- | --- | --- | --- | --- | --- |
|  | Pan02 | EMT6 | LL2 | 4T1 | CT26WT | RENCA | MC38 | MMTV-PyMT | B16F10 | GL261 |
| MNPs | 37.4 | 51.2 | 65.4 | 59.1 | 31.3 | 65.1 | 62.1 | 48.2 | 30.3 | 30.5 |
| Neutrophils | 19.4 | 16.3 | 9.9 | 7.0 | 4.1 | 4.0 | 2.6 | 2.0 | 0.5 | 0.1 |

(B)

| Cluster | % Neutrophils |
| --- | --- |
| G5a | 12.2 |
| G5b | 16.2 |
| G5c | 14.4 |
| Suppressive | 19.8 |
| Immature | 37.4 |

(C)

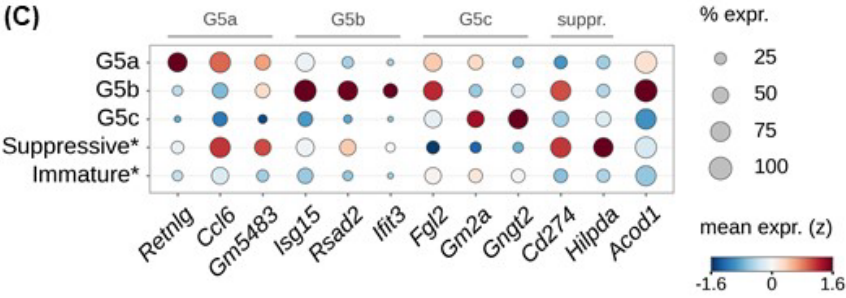

(D)

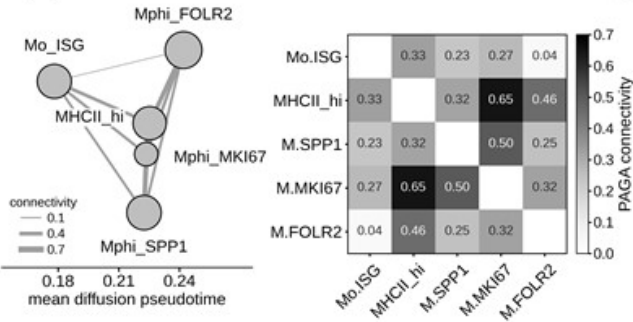

120,374 Mono/Mac cells, 10 models. Mono/Mac-specific PCA with the cell-cycle axis removed (max |r| of any PC with S/G2M score: 0.59 before, 0.00 after). Harmony over 30 samples. Node area = cell number; root = most monocyte-like Mo\_ISG cell.

(E)

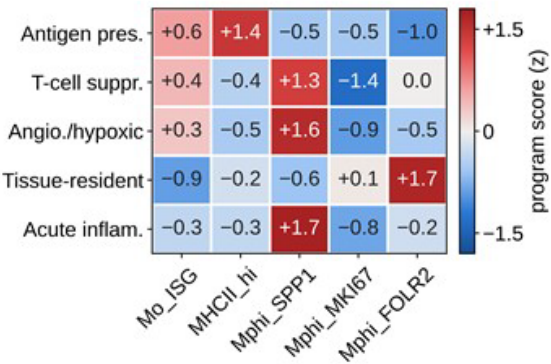

96,302 depth-matched Mono/Mac cells, 10 models. Each gene is z-scored across the five states first, then averaged, so every gene carries one vote.

(F)

| MNPs (10 models) | Spearman $\rho$ | P-value |
| --- | --- | --- |
| Mo_ISG | +0.60 | 0.073 |
| MHCII_hi | +0.56 | 0.096 |
| Mphi_MKI67 | -0.83 | 0.005 |
| Mphi_SPP1 | -0.13 | 0.73 |
| Mphi_FOLR2 | -0.60 | 0.073 |
| Mo_ISG+MHCII_hi | +0.70 | 0.031 |

(G)

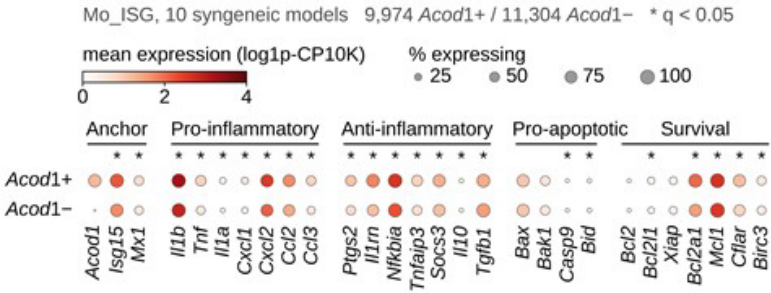
